# The prokaryotic origins of the COMMD protein family involved in eukaryotic membrane trafficking

**DOI:** 10.1101/2025.09.01.673571

**Authors:** Meihan Liu, Edmund R.R Moody, Farrah Blades, Caroline Puente-Lelievre, Kai-En Chen, Ella J. Stephens, Katharine A. Michie, Rosemary J. Cater, Tom A. Williams, Brett M. Collins, Michael D. Healy

**Author notes:** **Co-corresponding authors**: Brett Collins; Michael Healy.

## Abstract

The ten eukaryotic COMMD proteins are core components of the Commander complex, with central roles in endosomal membrane trafficking and signalling. Each protein has an α-helical N-terminal (HN) domain with a C-terminal copper metabolism gene MURR1 (COMM) domain. These ten family members assemble into a heterodecameric ring composed of five specific heterodimers. In this work we have combined structural homology searches with genome-wide predicted structures to identify ancestral COMMD-like proteins that exist as single genes in Bacteria and Archaea. Although there is limited sequence similarity to the eukaryotic proteins the bacterial and archaeal COMMD-like proteins are predicted to form homomeric ring-shaped assemblies like their eukaryotic counterparts. Our biophysical studies, crystal and cryo-EM structures confirm COMMD-like proteins readily form homooligomeric rings composed of eight or ten subunits assembled from core dimeric building blocks and inter-dimer interactions that are analogous to the heterodecameric core structure of the eukaryotic Commander complex. Phylogenetic analyses using amino acid sequences and FoldSeek structural alphabet (3Di) infer that the closest identified relatives to the eukaryotic COMMD proteins are found in Myxococcota bacteria. These findings indicate that COMMD genes emerged early in eukaryotic evolution through multiple rounds of duplication from a single ancestral gene likely acquired from bacteria.

## INTRODUCTION

The trafficking of transmembrane proteins and lipids through the endolysosomal system is critical to all eukaryotic life. The proteins and protein complexes that control this transport are generally conserved across the eukaryote domain from single celled protists to multicellular plants and animals^1^. Eukaryotes evolved from archaeal ancestors that formed a symbiotic partnership with a proto-mitochondrial bacterium^2,3^. The discovery of the Asgard archaea prompted further refinement of this model, with the Asgard lineage possessing numerous eukaryotic signature genes (ESGs)^4,5^, including many that encode proteins related to eukaryotic membrane trafficking and cytoskeletal organisation; such as the ubiquitin-ESCRT machinery^6^, RAB and ARF-like small GTPases^7,8^, SNAREs involved in membrane fusion^9^, as well as microtubules, actin and profilins^10–12^.

As eukaryotic endosomes mature to lysosomes, several large protein complexes direct the recycling of transmembrane proteins to other compartments such as the Golgi or plasma membrane to avoid degradation^13–15^. One such complex is Commander, made up of two sub-assemblies called the COMMD/CCDC22/CCDC93 (CCC) complex and the Retriever complex, the latter of which is structurally related to the endosomal Retromer machinery^16–18^. Mutations in multiple Commander subunits are causative of X-linked intellectual disability (XLID) and Ritscher-Schinzel syndrome (RSS), a multi-system developmental disorder characterised by abnormal craniofacial features, cerebellar hypoplasia, and stunted cardiovascular development^19^. Commander is composed of sixteen subunits divided into two subassemblies: the Retriever subassembly, which is composed of VPS29, VPS35L and VPS26C, and the CCC sub-assembly, which is comprised of ten copper metabolism gene MuRR1 domain (COMMD) proteins, two coil-coiled domain containing (CCDC) proteins CCDC22 and CCDC93 and a differentially expressed in neoplastic versus normal cells (DENN) domain protein DENND10. Together these sixteen components form a stably assembled complex that is localised to endosomal membranes, regulates transmembrane cargo transport.

At the heart of the Commander complex is a heterodecameric structure containing a single copy of each of the ten COMMD proteins COMMD1 to COMMD10^16–18^. A notable feature of this complex is that the ten proteins form a closed pseudo-symmetric ring, which is itself composed of five specific heterodimers. This ring is formed due to the unique structural features of the COMMD proteins which are composed of two domains; a helical N-terminal domain (HN domain) with a high degree of sequence divergence, and a more highly conserved C-terminal COMM domain. This COMM domain contains three β-strands followed by an α-helix, the unique arrangement of these elements leads to the formation of a hydrophobic core that results in the formation of an obligate dimer^20^. The decameric ring is then assembled from five of these dimers, where the COMM domains form a closed ring via β-sheet interactions between adjacent dimers. The N-terminal HN domains then extend peripherally on each side of this central core.

A number of studies have used amino acid sequence-based approaches to study the evolution of the Commander protein machinery^18,21–25^. All sixteen subunits are broadly conserved across the eukaryotic domain, including animals and their unicellular relatives such as choanoflagellates and amoebozoans. The complex has also been lost secondarily in some lineages, including in fungi^24^ (such as yeast) and flowering plants. A conservation pattern strongly suggesting that Commander subunits have co-evolved alongside the endosomal actin-nucleating Wiskott-Aldrich syndrome protein and SCAR homolog (WASH) complex^26^. Previous analyses have shown that all ten COMMD genes were likely present in the last eukaryotic common ancestor (LECA)^18^, with the series of gene duplications giving rise to each subfamily likely occurring along the eukaryotic stem (that is, prior to LECA). While search methods based on sequence similarity (such as BLAST and HMMer) did not find homologues of COMMD in prokaryotes^18,24,26^, a recent preprint used structure-based methods to identified COMMD-like proteins in the Asgard archaea, the closest prokaryotic relatives of eukaryotes^27^.

Here we use sequence and structure-based homology searches, evolutionary analyses, and biophysical assays to identify the origins and investigate the structure and function of prokaryotic COMMD-like proteins. We find that while CCDC22, CCDC93, DENND10, VPS26C and VPS35L are restricted to the eukaryotic lineage, ancestral COMMD proteins are present in both Bacteria and Archaea. These prokaryotic COMMD-like proteins have very low sequence identity (<20%) to any eukaryotic COMMD family members but share a conserved and highly similar fold including both HN and COMM domains. Phylogenetic analyses further indicate a close evolutionary relationship between eukaryotic COMMD proteins and a clade of bacterial sequences largely comprising representatives from the Myxococcota, bacteria that also appear to have contributed some other membrane associated genes to the nascent eukaryotic lineage^28^. We purified a selected set of bacterial and archaeal COMMD-like proteins and used mass photometry, X-ray crystallography and cryo-electron microscopy (cryo-EM) to demonstrate that most of these assemble into analogous dimeric structures to the human COMMD proteins, that then form oligomeric rings consisting of eight to ten subunits, which are homo-oligomeric rather than hetero-oligomeric as in eukaryotes. Our results indicate that the COMMD family of proteins evolved from a single prokaryotic ancestral protein-coding gene sharing key structural features, consistent with the early presence of endomembrane components during eukaryogenesis.

## RESULTS

### Ancestral COMMD-like proteins are found in Bacteria and Archaea

Previous work by us and others identified clear orthologues of each of the sixteen components of Commander in many eukaryotes, including choanoflagellates such as *Salpingoeca rosetta*^18,21,24,26^. To explore whether more distant structural homologues could be found in prokaryotes we used FoldSeek^29^ which allows for fast and accurate matching of structurally similar proteins, including structures of proteins encoded by bacteria and archaea available in the AlphaFold2 database^30–32^. These searches were performed initially using structural templates of Commander subunits from a variety of eukaryotic species, and then iteratively searching for additional prokaryotic proteins using identified hits (**Fig. 1A**). Using this approach, we confirmed the presence of clear orthologues of all sixteen subunits throughout the eukaryotic lineage and identified several previously unidentified homologues in *Reticulomyxa filosa* and *Anaeramoeba flamelloide –* single-celled organisms that are only distantly related to model eukaryotes (**Fig. 1B**). Turning to prokaryotes we could not identify any archaeal or bacterial proteins with significant structural similarity to CCDC22, CCDC93, VPS26C or VPS35L. Proteins with folds related to VPS29 (the shared subunit between Retriever and Retromer complexes) and DENND10 were identified in Archaea and Bacteria. Due to the abundant distribution of the metallophosphatase and DENN domain folds across all the tree of life we have not pursued these further. Using FoldSeek we were able to identify clear structural homologues of the COMMD proteins (**Fig. 1B and C**) that we have termed COMMD-like proteins. They have less than 20% sequence identity to any eukaryotic COMMD family members, explaining why they were not readily detected using sequence-based methods.

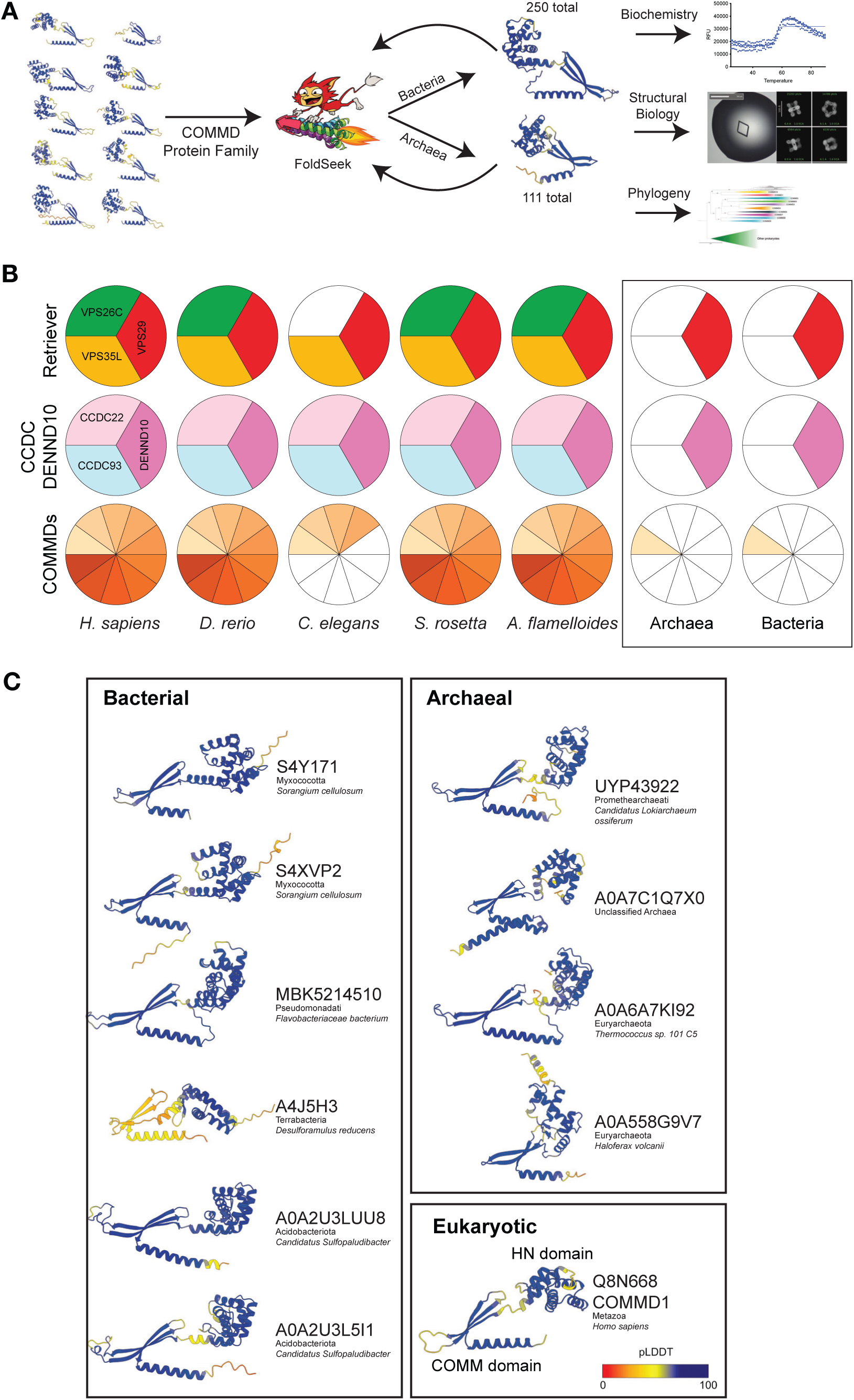

From initial searches using the ten human COMMD proteins against the UniRef100 AlphaFold2 database we identified 8,238 eukaryotic proteins with a COMMD-related protein fold. These were then used to screen the predicted proteins of all available prokaryotic genomes, with identified homologues used in subsequent iterative searches. In total we identified 250 bacterial COMMD-like proteins and 111 archaeal COMMD-like proteins broadly distributed across the diversity of prokaryotes (**Supplementary Dataset S1**). From Bacteria this included sequences from both Gracilicutes and Terrabacteria, including Proteobacteria, Chlamydia, Cyanobacteria and Actinobacteria amongst others. In the case of Archaea, we find examples across the evolutionary tree including in Methanobacteriota, Halobacteriales, Thermococcus and Asgard species. As discussed below, most prokaryotic genomes encode at most a single COMMD-like gene, with the exception of *Candidatus Sulfopaludibacter sp. SbA4* and a few species from the *Myxococcus* clade that encode two. A comparison of human COMMD1 and examples of COMMD-like proteins is shown **Fig. 1C**. The ten examples shown from Bacteria and Archaea are all predicted to possess N-terminal HN domains that shares the same α-helical topology as their eukaryotic homologues, and a C-terminal COMM domain comprised of a three-stranded anti-parallel β-sheet and extended α-helix.

### Structures and oligomeric properties of the prokaryotic COMMD-like proteins

We next characterised the structure of the ten exemplar COMMD-like proteins shown in **Fig. 1C** using both computational modelling and experimental approaches. A defining feature of the COMMD proteins in eukaryotes is that they form oligomeric complexes. The structure of the COMM domain requires the formation of an obligate dimer to bury their hydrophobic β-sheets^20^, the five specific heterodimers then assemble into a closed ring structure of ten subunits^16–18^. Although the eukaryotic COMMD proteins form heteromeric complexes *in vivo*, they are able to form homomeric assemblies when recombinantly expressed individually^20^. We first tested the propensity of the prokaryotic COMMD-like proteins to form homodimers using AlphaFold2 and found that they were universally predicted to form homodimers with high confidence metrics (**Figs. S1**-**S3**). We then modelled each of the ten COMMD-like proteins as increasingly larger oligomers (again using AlphaFold2), focusing on the octameric and decameric assemblies. Apart from A4J5H3 and A0A558G9V7 (for the sake of precise identification proteins have been labelled by either a Uniprot ID or Uniparc ID if Uniprot ID was unavailable) all proteins are predicted, with high confidence, to form closed ring structures using either eight or ten subunits (**Figs. S1**-**S3**). The predicted ring structures are assembled via similar interfaces between dimers and are structurally analogous to the known heterodecameric structures of the human COMMD1-10 complex^16–18^. A0A558G9V7 is predicted to form a decameric complex, however the geometry of its COMM domain does not allow for the octameric ring to close. A4J5H3 is not predicted to form a complex however it is worth noting that less than 10 sequences were identified which likely limits the ability of AlphaFold2 to make accurate predictions. These results suggest that the prokaryotic COMMD-like proteins may assemble into structures like their eukaryotic homologues.

We next aimed to experimentally validate the structures of these proteins and their oligomeric properties. Each of the ten proteins were expressed in *E. coli* and purified using His-tag affinity chromatography. By size exclusion chromatography all proteins have relatively homogenous elution peaks and Coomassie stained SDS-PAGE gels show that these proteins can be isolated with purity sufficient to obtain macromolecular crystals (**Fig. 2A** and **Fig. S4**).

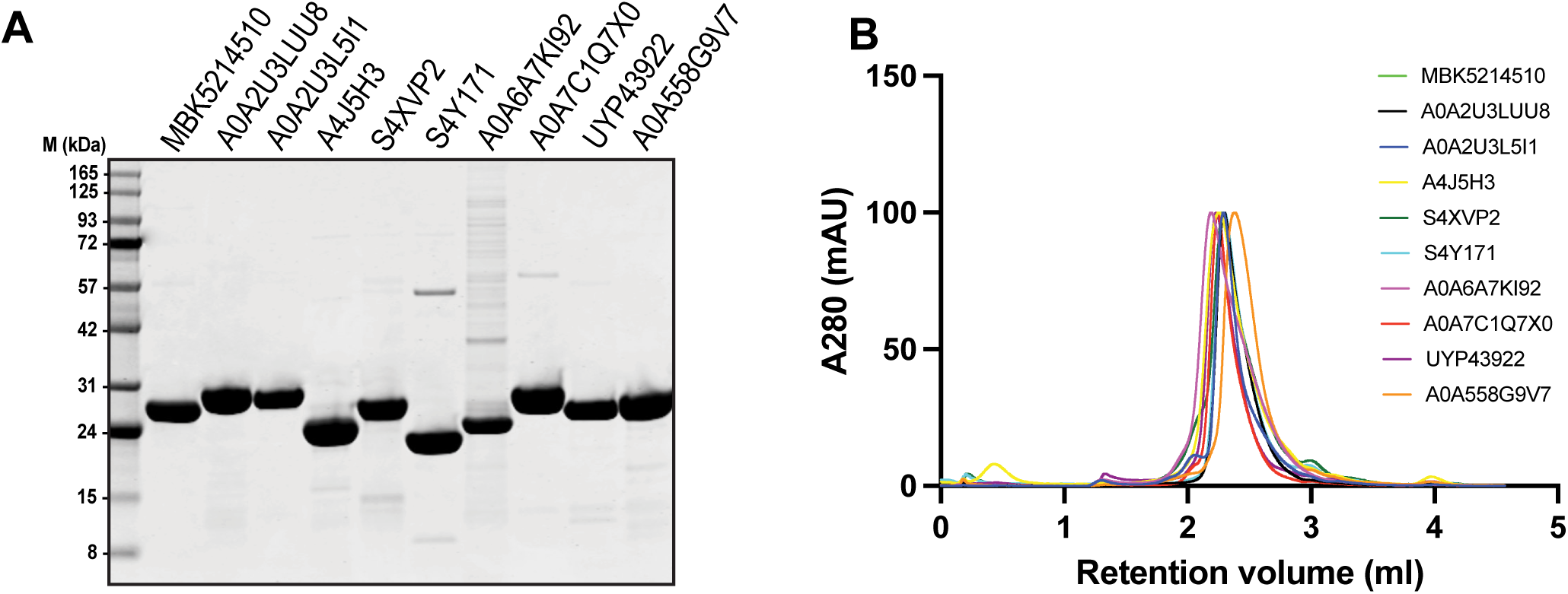

The retention volumes of the COMMD-like proteins by gel filtration are larger than expected for monomeric species suggesting they exist in homo-oligomeric complexes (**Fig. 2B**). To more precisely characterise these, we used multiangle laser light scattering (MALLS). Based on AlphaFold predictions and what is known about eukaryotic COMM domains, it is expected that all proteins form complexes that are composed of a dimeric building block. As indicated by MALLS, seven COMMD-like proteins form large complexes that consist of at least eight protomers based on the expected monomeric molecular weight (**Fig 3A; Figs. S5A and C**). The rapid decrease in molecular weight across the MALLS peaks for most proteins suggests that they are not fully monodisperse. To explore this further we used an orthogonal technique called mass photometry, which can provide an estimate of the heterogeneity within a sample and the molecular mass of each species (**Fig 3B-3D; Fig. S5B**). As an example, MBK5214510 shows evidence for the presence of dimeric, tetrameric, hexameric, octameric and decameric species. The formation of the oligomeric structure may be dependent on protein concentration (note MALLs in performed at ∼100x higher concentrations). A similar distribution of different oligomeric states can also be seen with A0A2U3LUU8, A0A2U3L5I1 and A4J5H3 (**Fig. S5B**). S4XVP2, S4Y171 A0A7C1Q7X0, UYP43922 and A0A558G9V7 form stable octamers, whereas A0A6A7KI92, seemed to form relatively stable decameric assembly. (**Fig 3C; Fig. S5B**). Interestingly, the distribution of oligomeric states is only seen in some of the bacterial homologues while archaeal homologues appear to be less dependent on concentration and tend towards existing in more stable oligomeric states as a result. As an exception the *myxococcota* (bacterial) homologues S4XVP2 and S4Y171 form stable homogenous octameric complexes in both MALLS and Mass photometry. One outstanding question is why we observe this variation in oligomeric assemblies in bacterial COMMD-like proteins. Broadly speaking there are two likely reasons: (i) the concentration at which the mass photometry measurements was taken may be too low for the larger oligomers to remain stably assembled, and/or (ii) additional components could be required to stabilise the assemblies as seen with the human COMMD proteins within the Commander complex^18^. To examine the stability of the COMMD-like protein oligomers, nano-DSF was used to determine the T_m_ ^50^ (the temperature at which 50% of the protein was unfolded). Unfortunately, several of the proteins had low, or no tryptophan residues required for fluorescence measurement which limited the assessment to five out of the ten. Of these, tested three were highly stable, with melting temperatures of ∼60°C and two were less stable having melting temperatures of ∼40°C. This did not appear to correlate with the oligomeric state of the COMMD-like protein or prokaryotic lineage (**Fig. S5D**).

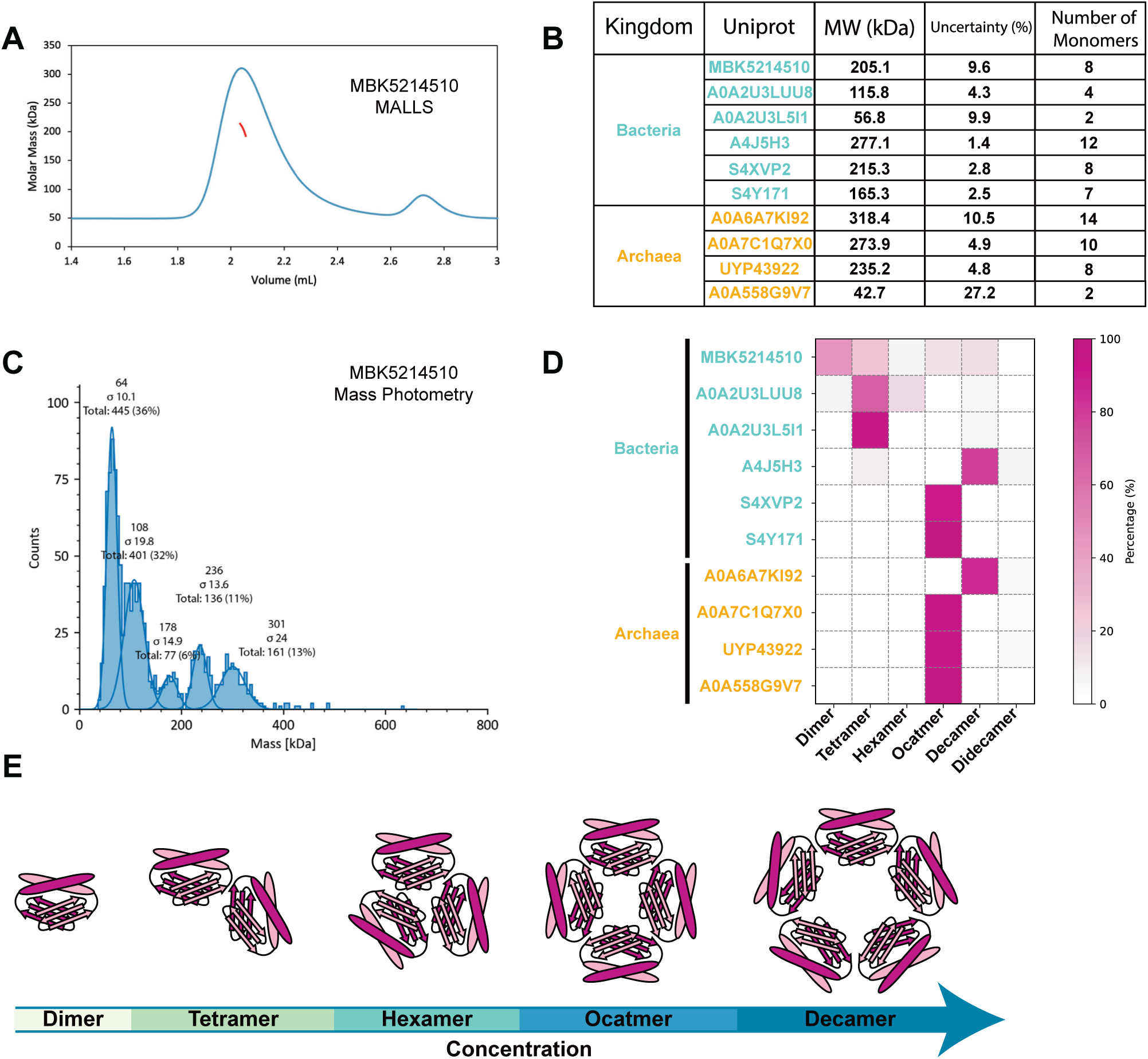

### Prokaryotic COMMD-like proteins form analogous complexes to the COMMD family

We next used Xray crystallography and cryo-EM structure determination to experimentally confirm our predicted structures of these COMMD-like proteins. Crystallisation screens of all ten COMMD-like proteins resulted in successful crystal growth, however most showed poor X-ray diffraction apart from crystals of MBK5214510 from *Flavobacteriaceae bacterium* (**Fig. 4A; Table S1; Supplementary Dataset S2**). We were able to solve the crystal structure at 2.6 Å resolution by molecular replacement using the AlphaFold2 model as a template. The asymmetric unit contains a dimer of MBK5214510 that resembles the human homo and heterodimeric structures solved previously^16,18,20^ (**Fig. 4A**). As expected, the dimer is formed by the intertwining of the two C-terminal COMM domains, each comprised of three β-strands and one α-helix that creates a large hydrophobic surface buried within the dimer interface. The HN domain contains six α-helices that form a tightly packed globular domain, the orientation with respect to the core COMM domain dimer results in the HN domains been arranged opposite and at 180° from each other (**Fig. 4A**). Inspection of the crystallographic lattice shows that the MBK5214510 dimer is forming an octameric ring structure along the 4-fold symmetry axis that is the same as the AlphaFold2 predicted octamer (**Fig. 4A**).

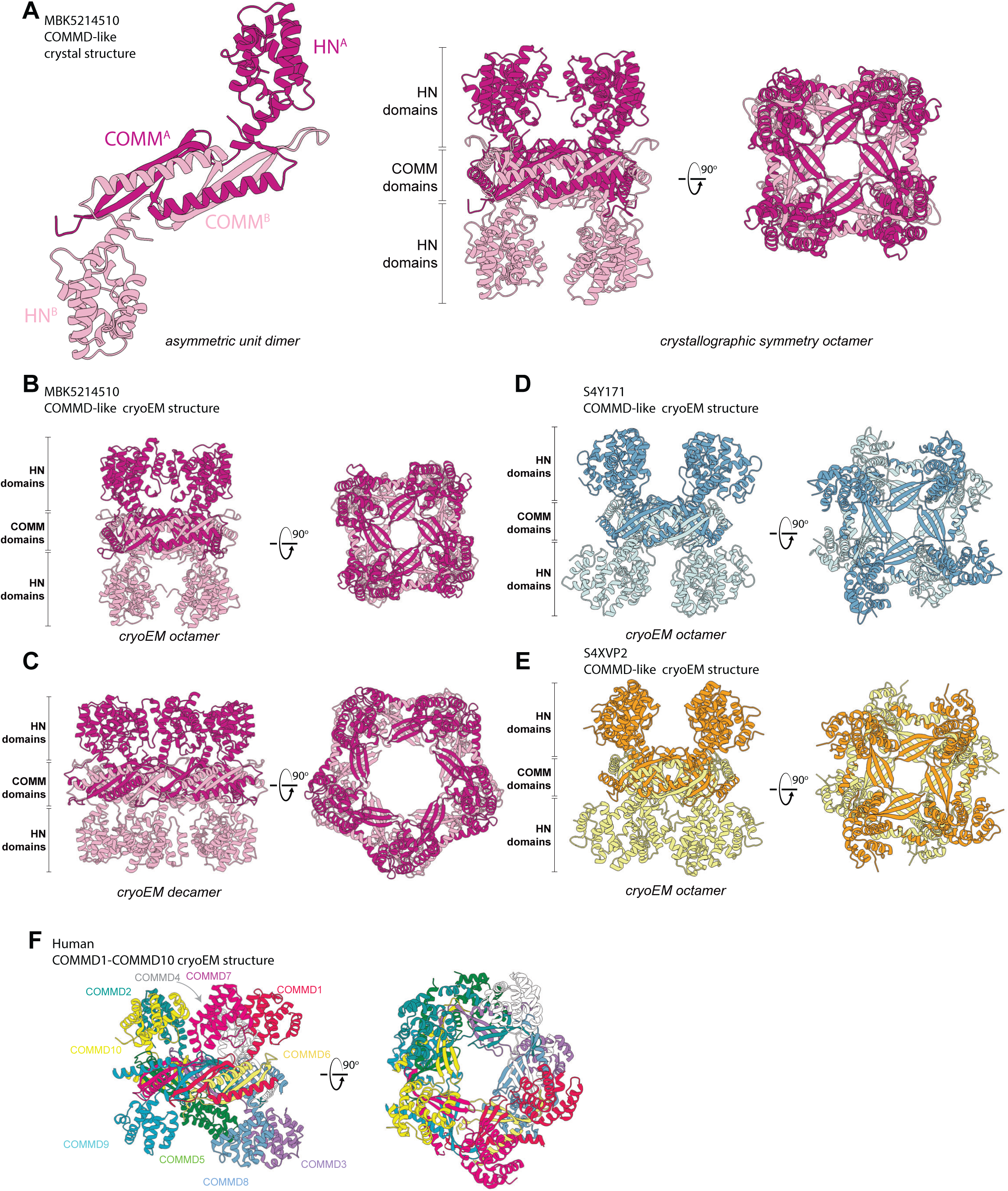

In parallel to the crystallographic studies, we also determined the structure of MBK5214510 by cryo-EM and single particle analysis (**Fig. 4B-4C; Fig. S6C-S6D; S7; Table S2**). Within the 2D classes there were clear examples of top views of both octameric and decameric ring structures. Filtering of these two different classes allowed us to resolve the octameric and decameric structures at ∼4.2 Å and ∼4.8 Å resolution respectively (**Fig 4B-4C; Fig. S7**). The octameric particles resembled the octameric ring we observed in the crystal structure, while the decameric particles were assembled in the same way as the octameric rings, but were composed of five identical homodimers rather than four. As mentioned above, AlphaFold2 predictions were able to produce confident models of closed ring structures using either 8 or 10 monomers (predictions using more or fewer subunits produce either fragments of rings (e.g A0A558G9V7) or physically unreasonable structures) (**Figs. S1-S3**). The modelling, biophysical data (**Fig. 3; Fig. S5**) and cryo-EM data (**Fig. 4B-4C**) thus supports the idea that the bacterial COMMD-like proteins have a capacity to form closed ring structures with some plasticity as to the number of dimeric building blocks that are incorporated. Comparisons of the experimental MBK5214510 oligomers to the AlphaFold2 predictions shows that the octameric crystal structure has the closest structural alignment with the predicted octamer (**Fig. S9**). The octamer determined by cryo-EM is highly similar to the prediction but shows greater flexibility than in the crystal structure. The decamer determined by cryo-EM is more divergent from the predicted decamer, with a significantly larger central pore, and is also wider than the cryo-EM structure of the human COMMD1-10 heterodecameric complex^17,18^ (**Fig. S9-S10**). Overall, even though there is only an average sequence conservation of less than 13% these results show that bacterial COMMD-like proteins are able to form homomeric rings that are analogous to the eukaryotic heterodecameric complexes, formed by a central ring of interlocked COMM domains that tessellate together and HN domains that extend peripherally.

To confirm that the octameric structure was conserved across species we also investigated those from the *myxococcota* - *sorangium cellulosum so0157 2*. This species contained two unique sequences (S4Y171 and S4XVP2) unlike many other species we identified. Using Cryo-electron microscopy we confirmed that these proteins also form octameric structures (**Fig. 4D-4E; Fig. S6E-S6F; Fig. S8; Table S2**) with a remarkably conserved architecture to both MBK5214510 and the human heterodecameric COMMD ring (**Fig. 4F**).

In the case of A0A6A7KI92 a low resolution cryo-EM dataset was collected that shows 2D classes consistent with an ability to form a homodecameric ring. Orientation bias prevented us resolving a high resolution structure (**Fig. S6**).

### Eukaryotic COMMD proteins are most closely related to COMMD-like proteins from the Myxococcota lineage

The evidence presented here suggests that the eukaryotic COMMD proteins originated via duplication and expansion of a single ancestral protein-coding gene. To assess the phylogenetic relationships between the Eukaryotic COMMD and the prokaryotic COMMD-like proteins, we performed independent phylogenetic analyses using amino acid sequences and 3Di structure data (**Figs. 5 and S11A**). These analyses did not support a close relationship between eukaryotic COMMDs and the COMMD-like proteins of Asgard archaea (the closest relatives of the eukaryotic nuclear lineage) or of the alphaproteobacterial relatives of the mitochondrion (**Fig. 5**), indicating that eukaryotes did not inherit COMMD from either of their immediate prokaryotic ancestors. However, robustly establishing the closest prokaryotic clade – that is, the extant relatives of the prokaryotic donor lineage - proved challenging because of the low level of similarity between sequences and substantial phylogenetic uncertainty in both sequence- and structure-based analyses, which were performed under the best-fit site-heterogeneous model (sequence analysis), and using a recently developed substitution model for 3Di characters (structure analysis^33,34^). Intriguingly, in the maximum likelihood sequence-based tree, eukaryotes branch with a clade predominantly comprising Myxococcota, but also including individual sequences from Desulfobacterota, Gemmatimonadota and Planctomycetota (bootstrap support 98%). This is intriguing because several other genes involved in eukaryotic membrane biology, including genes for sterol biosynthesis, appear to have been acquired from this clade^35^. A close and highly-supported relationship between eukaryotic COMMD and Myxococcota was also observed in the structure-based tree (**Fig. S11A)**.

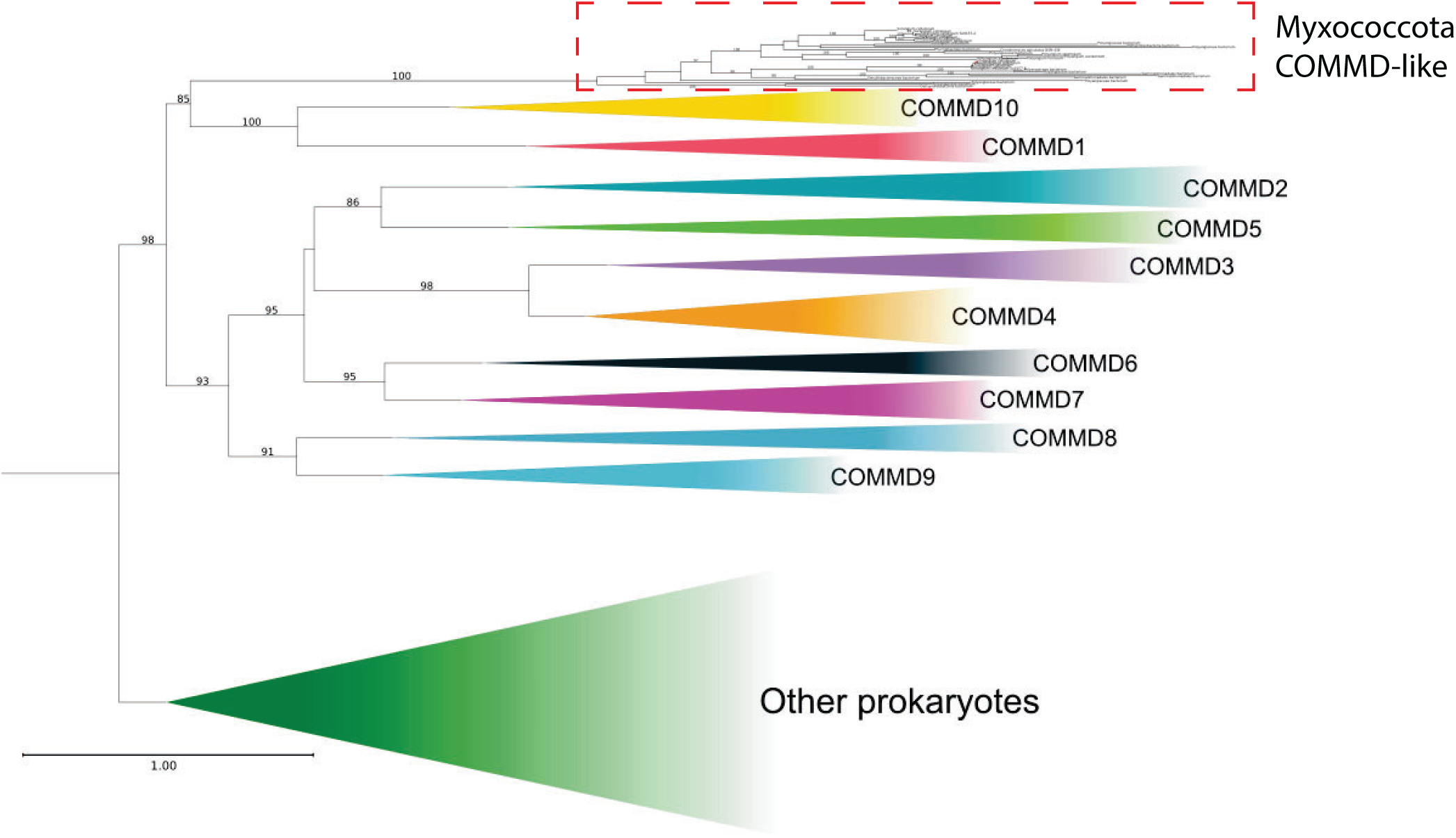

### *Myxococcota* COMMD-like genes cluster with secretion system components

Although the prokaryotic COMMD-like proteins clearly share a conserved structure with their eukaryotic relatives, their functions in either Bacteria or Archaea are unknown. In prokaryotes, genes that function in related pathways often tend to cluster together, for example components of hydrogenase complexes^36^ and the ancestral ESCRT and ubiquitination machinery in Archaea^37–39^. We therefore investigated the genomic context of the COMMD-like genes with a particular emphasis on the *Myxococcota* bacteria. An example genomic region focused on S4XVP2 from *Sorangium cellulosum* So0157-2 is shown in **Fig. 6A**. Interestingly, by investigating the adjacent genes, we noticed similarities between this region and the VgrG1B region of *Pseudomonas aeruginosa* PA01^40^ (**Fig. 6A**), although we note *P. aeruginosa* PA01 does not have an identified COMMD-like protein in the genome. The VgrG1B region is known to be critical for *P. aeruginosa* PA01 virulence and contains numerous components related to the Type VI secretions system (T6SS) (**Fig. 6B**).

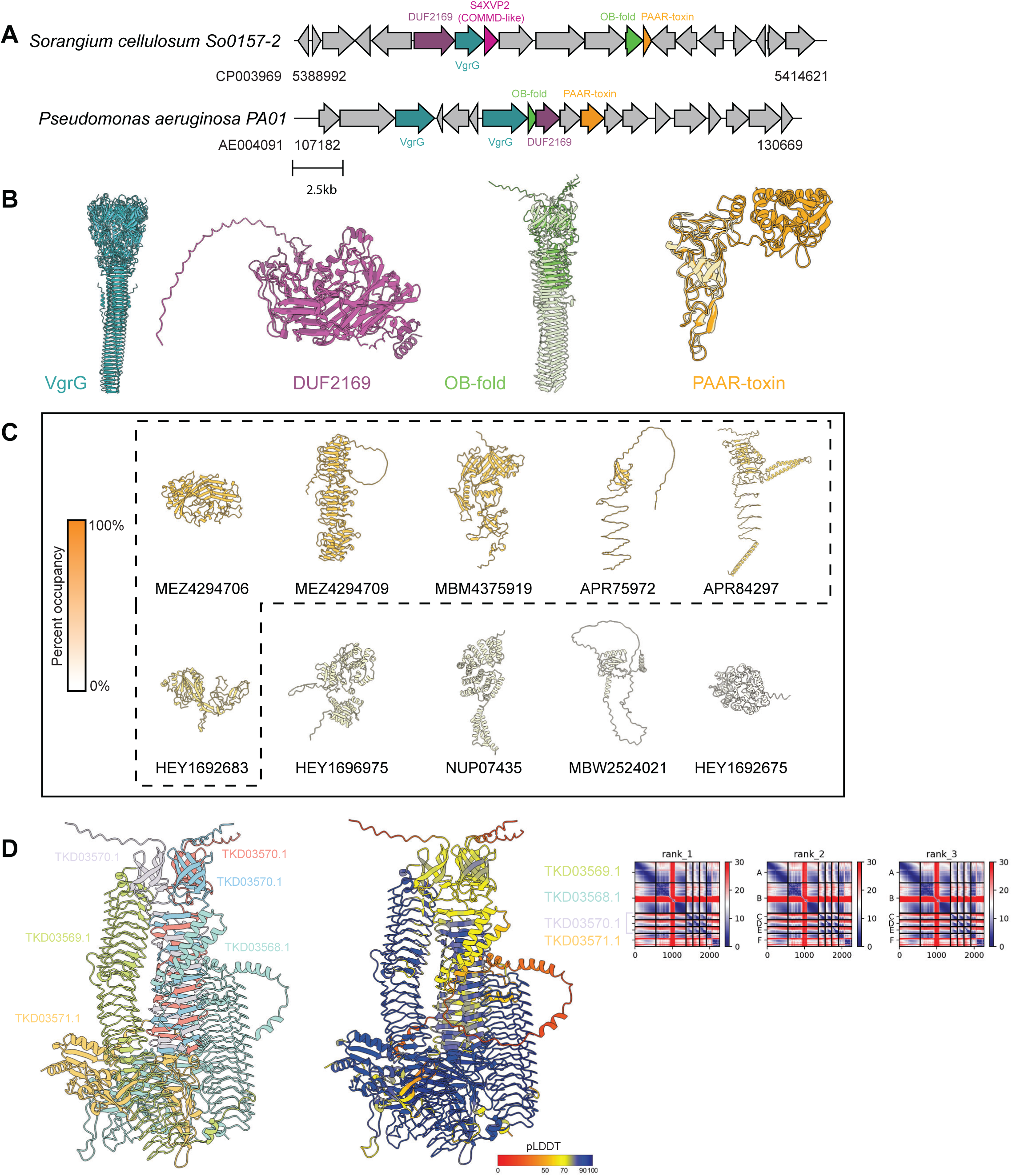

To understand if this was an isolated case or common to other COMMD-like proteins from *Myxcoccotta* we next performed syntenic analyses. Several COMMD-like homologues from *Myxococcota* species are available in the NCBI database that have not yet been catalogued in Uniprot or the AlphaFold2 database. In total, 55 full-length COMMD-like sequences were identified (compared to 32 from Uniprot and 92 in NCBI with shorter reads) with a surrounding region of at least 20,000 bp available (+/-10,000 bp). This region was exported and a clustered alignment with CLINKER^41^ was performed. This gave some hints of functionally related gene clustering at the sequence level (**Fig. S12**), but potential relationships became much more apparent at the level of predicted tertiary (and quaternary) structures. We then used AlphaFold2 to predict the structure of all the proteins in the extracted genomic regions (∼700 proteins) and the FoldSeek “easy-cluster” module to cluster these proteins^34^. From the ∼700 proteins 216 “clusters” were identified, of which 17 contained proteins that appeared within ± 10,000 bp of the COMMD-like gene at least 10% of the time. **Fig. 6C** shows an example protein structure from each of the top ten structural clusters coloured by the occupancy in relation to the 55 COMMD-like proteins. While the functions of many of these proteins are uncharacterised, those with the highest correlation are known virulence factors including OB-fold proteins (60%), VRG1 proteins (60%) and PAAR domain (47%) proteins. As mentioned above this region resembles the *P. aeruginosa* vgrG1b gene cluster and contains a protein with the DUF2169 fold (63%) (**Fig. 6B**). For these proteins and those with less clear structural identities FoldSeek was used to search the protein data bank for any related “ground truth” structures (**Table S4**). In addition to those proteins already mentioned this included a Serine-Threonine kinase^42^ (PDB: 4EQM), components of the *Mycobacterium tuberculosis* initial transcription complex^43^ (PDB: 6KON) attached to a tetratricopeptide repeat protein^44^ (PDB: 2PL2), HasB (PDB: 2M2K) and a thermal stable Old Yellow Enzyme^45^ (PDB: 3KRU).

Lastly, we performed iterative multimeric AlphaFold predictions of the proteins within the COMMD-like gene cluster to determine if COMMD-like proteins potentially interact with any of the proteins that were encoded nearby, but were unable to identify any putative COMMD-like-containing complexes (not shown). However, we did observe predicted interactions among several proteins that appeared to be structurally related to bacterial Type VI secretion system (T6SS) components. An example of such a complex from *Polyangium fumosum* is shown in **Fig. 6D**, composed of a homotrimer of TKD03570 and single copies each of TKD03571, TKD03569, TKD03568. The complex shares strong similarities to components of the T6SS baseplate machinery^46^.

### Other COMMD related proteins in Chlamydia, E. coli and Halobacteria

In addition to the Myxococcota bacteria, we also noted an expansion of COMMD-like proteins in *Chlamydia* and *Halobacteria*. The exact reason for this expansion is unclear, and a potential caveat is that due to their respective importance in disease and Archaeal evolution respectively, the *Chlamydia* and *Halobacteria* clades have many representative genome sequences compared with other species we identified. It is also worth noting that while the overall structure of the *Chlamydia* proteins conforms to the classical COMMD-fold, containing a helical HN domain and a COMM domain, they appear to function in a unique manner. This is seen in two previously determined crystal structures (PDB: 3Q9D and 4MLK^47^) in which they form a hexameric assembly coordinated by interactions between the HN domain as opposed to the classical β-sheet augmentation seen in the COMMD and COMMD-like structures presented here and elsewhere. Regardless we investigated the genomic context of these proteins to see if there was continuity with *Myxococcota*. This showed that these COMMD-like proteins clearly exist within species level clusters (**Fig S12-14**). In **Table S5** the clusters are summarised for an exemplar COMMD-like protein: O84588 for *Chlamydia trachomatis serovar D (strain ATCC VR-885 / DSM 19411 / UW-3/Cx)* and G0HSR7 for *Haloarcula hispanica (strain ATCC 33960 / DSM 4426 / JCM 8911 / NBRC 102182 / NCIMB 2187 / VKM B-1755)*. As before a combination of AlphaFold2 and FoldSeek was used to understand the type of genes these cluster with and if any complexes were forming with nearby encoded proteins. In both cases the COMMD-like protein was not predicted by AlphaFold to make direct interactions with any of the proteins within the respective gene cluster.

## DISCUSSION

This work identifies archaeal and bacterial proteins that are structurally homologous to the eukaryotic COMMD protein family. Our analyses show that many species of bacteria and archaea encode either one or two COMMD-like proteins compared to the ten distinct genes that are conserved across the eukaryotic lineage. Of note, a recent preprint also reported the existence of COMMD-like proteins in Asgard archaea, supporting these findings^27^. While the evolutionary path from these single prokaryotic COMMD-like genes to the expanded eukaryotic repertoire is not entirely clear, our phylogenetic analyses suggest that eukaryotes did not acquire COMMD genes directly from their Asgard or alphaproteobacterial ancestors, but likely from another bacterial lineage. Among extant bacteria, Myxococcota appear to encode the sequences most closely related to eukaryotic COMMD proteins, although this inference may be limited by the resolving power of current phylogenetic methods on distantly related sequences.

Our structural and biophysical analyses confirm that the prokaryotic COMMD-like proteins are driven to form homodimers using the same structural principles as the heterodimers formed by specific eukaryotic COMMD family members. Furthermore, these can then assemble into homo-oligomeric ring complexes that are structurally analogous to the hetero-decameric rings formed by the ten eukaryotic proteins. Several other protein families in prokaryotes have some structural similarities to the COMMD proteins. These include the CT584/Cpn0803 proteins that are exclusively found in Chlamydiales^47,48^, and the Whirly/Pur-α family of DNA binding proteins that are found in species across the three domains of life^49–54^. Despite superficial similarities, these are not strictly COMMD-like proteins. The CT584/Cpn0803 proteins have C-terminal domains that form dimers in a similar way to the COMM domain, and a helical N-terminal domain that has a related topology to the HN domain, but they assemble into very different hexamers that are unrelated to the homo and heterooligomeric rings formed by the COMMD protein family^47,48^. The Whirly/Pur-α proteins lack any structures related to the COMMD HN domain, but have small, repeated domains composed of a four-stranded β-sheet and a C-terminal α-helix that can form dimeric structures in a similar way to the COMM domain^49–54^. Unlike members of the COMMD or COMMD-like protein family Pur-α proteins have a highly charged surface allowing for DNA/RNA binding. Although it would appear COMMD-like proteins lack the surface residues required to directly bind DNA/RNA it is notable that in both Chlamydiales and Haloarchaea that genes for DNA modulation machinery are found in close proximity to the COMMD-like proteins. Altogether these proteins do not share any sequence homology, lack key structural elements and assemble in very different ways to the COMMD proteins, so it is unclear if they share an evolutionary history or are examples of convergent evolution to superficially similar structural folds.

Many different protein families have evolved the ability to form multimeric ring structures, which most commonly involves homomeric symmetric interactions with one or a few repeated protein subunits^55–58^. Heteromeric pseudo-symmetric structures composed of multiple paralogous proteins, as seen for the eukaryotic COMMD decamer^16–18^, are less common and typically arise from a series of gene duplication events converting homomeric into heteromeric complexes. Some prominent examples include the eukaryotic 20S proteosome^59^, the TRiC/CCT chaperonin complex^60–62^, the Sm/Lsm RNA-binding proteins^63–65^, the CMG helicase^66–68^ and the origin recognition complex^69,70^ involved in initiating DNA replication. In the TRiC/CCT chaperonin complex for example, there is ample evidence of these duplication events and subsequent sequence divergence that creates eukaryotic heteromeric assemblies with increasing complexity that are structurally analogous to the homomeric counterparts in archaeal species^71^. In the case of the Sm/Lsm rings, much like the eukaryotic COMMD family, there is a clear distinction between eukaryotes, which possess a full complement of fourteen Sm/Lsm genes, and prokaryotes that have only a single Sm/Lsm paralogue, suggesting the duplication events occurred very early in the eukaryotic branch. It has been argued that the recruitment into the complex spliceosomal machinery is at least partly responsible for creating such a large paralogous family in eukaryotes derived from the single gene precursors in Archaea and Bacteria^72,73^, and we similarly propose that the high degree of functional divergence and specialisation of the eukaryotic COMMD proteins from the prokaryotic COMMD-like proteins reflects their incorporation into the Commander and co-evolving WASH assemblies.

Unlike the COMMD proteins we did not uncover any prokaryotic proteins with clear similarity to the other subunits of Commander with the exception of VPS29 which has a typical metallophosphatase fold found in numerous proteins across the three domains of life and DENND10 which similarly contains the common DENN domain fold. The Commander complex is widely but not universally conserved in eukaryotes; for example, it has become secondarily simplified in some plants and is completely absent in fungi, although most eukaryotic lineages retain all ten subunits that date back to LECA^18^. During eukaryotic evolution, these subunits have co-evolved with the endosomal actin remodelling WASH complex^21,24,26^. Thus, in eukaryotes, the COMMD proteins appear to be essential core components of the Commander machinery that regulate endosomal trafficking and cytoskeletal organisation. By contrast, the function(s) of the bacterial and archaeal COMMD-like proteins is currently unclear.

The origin of eukaryotes involved symbiosis between an Asgard archaeon and an Alphaproteobacterium^74^. The bacterium evolved to become the mitochondria of modern eukaryotes, while the Asgard contributed key nuclear and cytosolic machinery including information processing (that is, DNA replication, transcription and translation), membrane trafficking and cytoskeletal organisation^6–9,11,12,37,75^. Although COMMD-like proteins are present in Asgards^27^, there do not appear to be any other components of the Commander or WASH machinery in Archaea that suggest a nascent role in membrane or cytoskeletal regulation. Indeed, phylogenetic analysis instead suggests that the eukaryotic COMMD family is most closely related to sequences from modern *Myxococcota*. Several other core eukaryotic genes with roles in membrane biology show phylogenetic affinities with *Myxococcota*, including key genes of sterol biosynthesis^35,74^. One possibility is that these were acquired by horizontal gene transfer during the origin of eukaryotes, potentially mediated by viruses^76^. An alternative explanation is provided by the syntrophy hypothesis for the origin of eukaryotes, in which the symbiotic interactions between Asgard and Alphaproteobacterium occurred within a delta-proteobacterial (potentially myxococcotal) host^28^. However, it is currently unclear whether an additional cell biological partner is needed to explain the acquisition of these genes from *Myxococcota* during early eukaryotic evolution, particularly given evidence for additional genetic contributions from other lineages^77,78^. We were unable to conclusively determine the function of the closest relatives of COMMD in modern *Myxococcota*, although our synteny analyses suggest a potential association with a Type VI related secretion system in the *Myxococcota* clade.

In summary, the combination of genome-wide structure prediction, and structural homology searches and phylogenetics have provided a powerful and comprehensive approach for identifying remote protein homologies that are difficult - if not impossible - using sequence-based methods alone. Eukaryotic COMMD proteins have a unique structural organisation that drives heteromeric assembly into stable ring-shaped complexes, and both archaea and bacteria possess ancestral COMMD-like proteins that form analogous complexes with only a single repeated subunit. This supports the idea that the eukaryotic genes have evolved from a single ancestral prokaryotic gene and have undergone repeated duplication and evolution to allow for a highly specific set of heteromeric interactions to occur. While the functions of the prokaryotic COMMD-like proteins remain obscure, and the evolutionary process that promoted COMMD duplication and co-evolution with other Commander subunits is currently unknown, these questions will present a rich avenue for future investigation on the origins of eukaryotes. Furthermore, studies of the simpler prokaryotic COMMD-like proteins may provide new insights into the functions of the eukaryotic COMMD assembly.

## METHODS

### Structure-based homology search

FoldSeek^29^, with the full UniProt database, was installed locally. The initial analysis utilized a folder containing the predicted structures of the following human proteins: COMMD1-10 (excluding COMMD6 to focus on full-length COMMD proteins), VPS26C, VPS29, VPS35L, CCDC22, CCDC93, and DENND10. This folder was used to search the UniProt database, restricted to eukaryotic sequences. Default settings were applied, except for coverage, which was set to 90% to exclude PUR-α and its homologues from the results. Proteins identified with a confidence score of 1 were downloaded from the AlphaFold database to create a new starting dataset. This updated folder was then used in a second screening of the UniProt database. The iterative screening process continued until the increase in the total number of identified proteins was less than 2%. The final datasets obtained from the eukaryotic screen were subsequently used as starting points for searches in the archaeal and bacterial databases. The same iterative method was applied to these datasets.

### Protein structure predictions

All individual proteins were obtained from the AlphaFold2 database^31^ unless no structure was available. In such cases structures were generated using a local instillation of the ColabFold^79^ v1.5.5 implementation of AlphaFold2-PTM^30^. A single model was used with 3 recycles to generate additional structures, quality of model was assessed by the ptm score. Protein complex predictions were generated using ColabFold v1.5.5 implementation of AlphaFold2-multimer^79,80^, 5 models were generated with 3 recycles. Model quality was determined by extent of overlap and iPTM score. All images were generated using chimeraX^81^.

### Molecular cloning

The sequences for 10 full-length COMMD-like proteins were synthesized and cloned for bacterial expression into the pET-28a(+) vector by Gene Universal (USA). The two different cloning sites are Ncol & Xhol (A0A558G9V7, A0A6A7KI92, A4J5H3, S4XVP2, S4Y171) and Ndel & Xhol (A0A2U3LUU8, A0A2U3L5I1, A0A7C1Q7X0, MBK5214510, UYP43922). S4XVP2 and S4Y171 were ordered as gene fragments and cloned into a modified version of the LM627 vector (addgene: 191551) containing GST using the Golden Gate cloning method as described by Wicky *et al*^82^. S4XVP2 and S4Y171 produced more homogeneous higher quality protein when expressed with a N-terminal GST tag compared with the C-terminal His tag.

### Protein expression and purification

All the proteins were expressed using *Escherichia coli* BL21(DE3) competent cells grown in the presence of kanamycin at 37°C. Protein expression was induced by 1 mM isopropyl β-D-thiogalactopyranoside (IPTG) when cell density OD600 reached 0.8, the temperature was then reduced to 25°C and cultures were incubated overnight. The cells were harvested the following day by centrifugation (Beckman JLA 8.1 rotor) at 7,000 xg, 4 °C for 10 minutes and resuspended in lysis buffer (500 mM NaCl, 50 mM Tris-HCl (pH 8.0), 5 mM imidazole, 10 % glycerol, 2mM β-mercaptoethanol, benzamidine, and DNAseI).

For protein purification, cells were firstly lysed using a Constant systems cell disruptor at 35 Kpsi, followed by centrifugation (Beckman JLA 16.250 rotor) at 20,000 xg, 4°C for 30 minutes, after which the soluble His-tagged proteins were contained in the supernatant. Subsequently, TALON resin (Bio-strategy Pty Limited) was added to the supernatant for binding His-tagged proteins and eluted 500 mM NaCl, 500 mM imidazole, 10% glycerol, 2 mM β-mercaptoethanol. Proteins were further purified using pre-equilibrated size exclusion chromatography (SEC) columns in 100 mM NaCl, 20 Tris-HCl (pH 8.0), either a Superdex 200 Increase 10/300 GL column (Cytiva) or HiLoad 16/600 Superdex 200 pg column (Cytiva) attached to an AKTA system (GE Healthcare). The purified protein fractions were checked by sodium dodecyl sulfate-polyacrylamide gel electrophoresis (SDS-PAGE) using Bolt 12 % Bis-Tris gel in 1x MES running buffer at 165 V for 45 minutes.

### Size exclusion chromatography-Multiangle laser light scattering (SEC-MALLS)

Each protein sample at 1 mg/ml (30 ul) was injected onto the Superose 6 increase 5/150 GL column equilibrated with the buffer containing 100 mM NaCl, 20 mM Tris-HCl (pH 8.0). MALS was performed using a DAWN HELEOS II 10-angle light-scattering detector paired with an Optilab rEX refractive index detector (Wyatt Technology), attached to a Prominence high-performance liquid chromatography (HPLC) (Shimadzu) at a flow rate of 0.25 ml/min.

### Single-Particle Mass Photometry

Molecular mass of all the proteins was measured by a Refeyn OneMP mass photometer (Refeyn Ltd) in the CMM. Initially bovine serum albumin (BSA) from the Sigma-Aldrich was applied as a mass calibration for MBK5214510, given MBK5214510 ability to for concentration dependent oligomers this was used in subsequent measures for calibration. For actual measurement, 10 μl of 100 nM protein samples (200 nM for A0A2U3L5I1, A0A558G9V7) was added to the coverslip. Movie were collected for 180 s. The data was then analysed using Aquire^MP^ software and generated a gaussian distributions to determine the protein mass composition.

### Nano-Differential Scanning Fluorimetry

NanoDSF was conducted using Prometheus NT.48 equipped with backscattering detectors (NanoTemper Technologies, München, Germany) at the Protein Expression Facility (PEF, The University of Queensland). Each protein sample was prepared in triplicate at a concentration of 1 mg/ml and loaded into individual NanoDSF grade high sensitivity capillaries (NanoTemper Technologies, München, Germany). The samples were then exposed at 100% excitation power, with a temperature gradient ranging from 20°C to 95°C with a ramping rate of 1°C/min. Tryptophan fluorescence emission was collected at 330 nm and 350 nm using a dual-UV detection, meanwhile, protein aggregation was monitored by backreflection optics. The PR.ThermControl software was used to calculate thermal stability parameters (Tonset, Tm, and Tagg).

### Crystallisation and structure determination

Crystallization trials were performed on a Mosquito liquid handling robot (TTP LabTech) in the commercially available SPARSE matrix 96-well hanging-drop plates (Index, LMB, MIDASplus, PEG/Ion, PEGRx and JCSG+) at 20 °C. Prior to sample loading into wells, protein samples (except S4XVP2, S4Y171) were concentrated to ∼10 mg/ml (determined by Bradfold). Crystals hits were observed for all proteins across different plates. To scale up the drop and crystal size, several conditions were optimized in a 24-well hanging drop vapor diffusion (Hampton) plate covered by glass cover slips. On each glass coverslip, three drops were prepared with protein and reservoir solution mixed in ratios of 1:1, 2:1 and 3:1 (protein:reservoir solution). Glycerol (10 %) was added into the mother liquor as cryo-protectant before flash-freezing in liquid nitrogen. The crystals used for MBK5214510 (15mg/ml) structure determination were grown in MIDASplus C9 (10 % v/v DMSO, 0.1M NaCl and 25% PE5/4 which in this case served as the cryo-protectant) a final concentration of 15mg/ml.

X-ray diffraction data were collected at the Australian synchrotron on the MX2 beamline. The data was indexed and integrated with XDS^83^ and scaled with AIMLESS withing the CCP4 suite^84^. The initial structure was resolved using molecular replacement in Phaser^85^ with the AlphaFold2 structure as an initial template. Further refinements of the structure were performed with PHENIX^86^, COOT^87^ and the chimera plugin ISOLDE^88^.

### Cryo-EM grid preparation and data collection

Freshly glow-discharged Quantifoil 1.2/1.3 Copper 200 mesh grids were used for cryo-EM data collection. For MBK5214510, A0A6A7KI92, S4XVP2 and S4Y171 a final concentration of 10 mg/ml in 100 mM NaCl, 20 Tris-HCl (pH 8.0) was loaded onto grids. Grid vitrification was performed in liquid ethane using a Vitrobot Mark IV plunge freezer (Thermo Fisher), using a 3 μL sample application, 95% humidity, 4°C, minus 1 blot force and 2.5 s blot time. After freezing, grids were stored immediately under liquid nitrogen until image acquisition.

Cryo-EM data for MBK5214510 was acquired using a JOEL CRYOARM 300 (JOEL) microscope operated at an acceleration voltage of 300 kV, equipped with an in column Ω energy filter (20 eV slit width) and a Gatan K3 direct electron detector. A total of 1,905 movies, at 60,000 x magnification, with a pixel size of 0.82 Å per pixel and a total exposure dose of 40 e/Å2, within a target defocus range from −2.5 to −0.5 μm. A0A6A7KI92 movies (769 movies) were collected on JOEL CRYOARM200 at 200 kV under 100,000 x magnification (corresponding to calibrated pixel sizes of 0.52 Å/pixel). This microscope included a cold-field emission gun (FEG), an in-column Ω energy filter and a Gatan K2 direct electron detector. As before the total exposure dose was maintained at 40 e/Å2, with the defocus range adjusted between −2.5 and −0.5 μm.

Cryo-TEM data for S4XVP2 and S4Y171 were collected on a Glacios 2 cryo-transmission electron microscope (Thermo Fisher Scientific) operated at an acceleration voltage of 200 kV and equipped with an integrated Thermo Scientific Falcon 4i Direct Electron Detector, an extreme high-brightness field emission gun (X-FEG), and Thermo Scientific EPU software. A total of 5,993 and 3,432 movies, respectively, were recorded at 190,000 x magnification, with a pixel size of 0.75 Å per pixel, a total electron exposure of 50 e/Å2, within a target defocus range from -2.5 to -0.5 μm.

### Cryo-EM data processing

Data processing of MBK5214510, A0A6A7KI92, S4Y171 and S4XVP2 was performed using the University of Queensland’s remote web access desktops (https://github.com/UQ-RCC/hpc-docs/blob/main/guides/OnDemand-Guide.md) in CryoSPARC V4.6.1^89^. For MBK5214510, 1,905 movies were imported and motion corrected using Patch Motion Correction, followed by CTF estimation. During the subsequent exposure curation step, the contrast transfer function (CTF) fit resolution was restricted to 6 Å, and the median relative ice thickness was adjusted to 1.031, resulting in the retention of 1,331 micrographs. From this dataset, 253,056 particles were initially blob-picked with a particle diameter range from 100 Å to 160 Å and template-picked at 160 Å. Following iterative rounds of 2D classification (**Table S2**), only the high-quality 2D classes were selected as templates for further template picking (150 Å particle diameter) and 2D classification (**Fig S7**). Ultimately, 120,876 particles were chosen and used to construct two ab-initio models. At this point, octameric and decameric structure volumes were separated and reconstructed using non-uniform (NU) refinement with D4 and D5 symmetry imposed. The final maps were obtained at resolutions of 4.22 Å for the octamer, and 4.77 Å for the decamer.

The 769 movies collected for A0A6A7KI92 were processed similarly as MBK5214510, with a particle diameter ranging from 120 Å to 200 Å for blob picking and 100 Å for template picking. As a result, 66,476 particles were selected from 2D classification. Due to a strong bias in orientations towards the top view, no further processing was attempted.

The 5,993 and 3,432 movies for S4Y171 and S4XVP2, respectively, were initially processed as MBK5214510. However after particle extraction the protocol varied, in both cases we used more stringent 2D classification and ab-initio model building following a recent protocol from Kim et al^90^. Following multiple rounds of 2D classification we selected a small subset of extremely high quality particles to build a high quality ab-initio model (**Fig S8**). This initial volume was then used in 2 rounds of heterogenous refinement with 4 junk classes generated from a prematurely terminated ab-initio job. These particle stacks were then refined using NU-refinement with an imposed D4 symmetry (**Fig S8**).

### Sequence-based phylogenetic analysis

Prokaryotic sequences from above were aligned using MAFFT v7.505^91^ (--linsi), this alignment was used to infer maximum likelihood trees using IQTREE 2.2.6^92^, using 20 profile mixture models (LG+C20)^93^ with empirical amino acid frequencies derived from the data (+F), four gamma categories (-G)^94^, with 10,000 ultrafast bootstraps^95^. Additional trees were inferred under the same settings, but with alternative constraints built using TreeViewer^96^, one constrained with the Myxococcota to be monophyletic with respect to other bacteria in the tree, the other constrained the archaeal sequences to be sister to the eukaryote COMMD proteins (to the exclusion of all bacterial sequences). These constrained topologies and the maximum likelihood topology were then compared using the approximately unbiased^97^ test under the same model, with 100,000 RELL replicates^98^. This analysis could not significantly reject (P < 0.05) the *Myxococcota* clade branching with other bacteria, or the archaeal sequences being more closely related to the eukaryotic COMMD sequences. However, imposing the archaeal constraint resulting in a subtending branch uniting eukaryotes and archaea of length 0.0000030 substitutions per site, suggesting the archaeal sequences are unlikely to be sister to the eukaryotic sequences. In all of these analyses, the Asgard sequences were resolved as only distantly related to the eukaryotic sequences, perhaps because of relatively recent horizontal acquisition from bacteria.

### Structure-based phylogenetic analysis

Protein structures were downloaded from the AlphaFold database (where available) or predicted using AlphaFold2. Structural alignment was performed using Foldmason^99^ and visualised with AliView^100^. Phylogenetic trees were inferred from the 3Di alignment with IQ-TREE v3.0.183^92^, 10,000 ultrafast bootstraps^95^, using the Foldseek 3Di matrix^29^. This is a fixed scoring matrix that was pre-trained with structurally aligned proteins to model the transition probabilities between structural 3Di states. Scores are based on observed versus expected pair frequencies using empirical background frequencies of 3Di states. Phylogenies were visualised and prepared for publication using iTOL v7.5^101^.

### Genomic context analysis

COMMD-like proteins were identified and enterz was used to download the region 10,000bp upstream and downstream of that reference point. CLINKER^102^ was then used to align these genomic regions based on sequence identity. In the case of Myxcoccota sequences they were extracted from the .gbk file and the structure of those sequences was predicted using AlphaFold2 as described above.

## DATA AVAILABILITY

The crystallographic and cryo-electron microscopy data generated in this study have been deposited in the PDB database under accession code 9Q6A [https://doi.org/10.2210/pdb9Q6A/pdb], 9Q6C [https://doi.org/10.2210/pdb9Q6C/pdb], 9Q6D [https://doi.org/10.2210/pdb9Q6D/pdb], 9ZA1 [https://doi.org/10.2210/pdb97A1/pdb] and 9ZA2 [https://doi.org/10.2210/pdb97A2/pdb]. In addition, the associated cryo-electron density maps have been deposited in the EMDB uder accession code EMD-72266 (9Q6C) [https://www.ebi.ac.uk/emdb/EMD-72266], EMD-72267 (9Q6D) [https://www.ebi.ac.uk/emdb/EMD-72267], EMD-73963 (9ZA1) [https://www.ebi.ac.uk/emdb/EMD-73963] and EMD-73964 (9ZA2) [https://www.ebi.ac.uk/emdb/EMD-73964]. The AlphaFold2 modelling data used in this study are available in the ModelArchive database under accession code ma-commd-like [https://modelarchive.org/doi/10.5452/ma-commd-like].

## ACKNOWLEDGEMENTS

We acknowledge use of the University of Queensland Remote Operation Crystallization and X-ray (UQ ROCX) Facility. X-ray data were collected on the MX1 and MX2 microfocus beamline at the Australian Synchrotron. The authors acknowledge the support of the Microscopy Australia Research Facility at the Centre for Microscopy and Microanalysis at The University of Queensland. We wish to acknowledge The University of Queensland’s Research Computing Centre (RCC) for its support in this research^103^. We would like to thank Prof. Paul Curmi for prompting this work with an insightful suggestion over drinks at the Lorne Protein meeting. To paraphrase, “Remember how during your PhD you thought it might be fun to look for homologues of your protein in archaea? You should try that again!”.

## FUNDING

B.M.C. is supported by an Investigator Grant from the National Health and Medical Research Council (APP2016410) and an Australian Research Council (ARC) Discovery Project Grant (DP240101315). M.D.H is supported by an Investigator Grant from the National Health and Medical Research Council (GNT2042760) and a project grant from Dementia Australia Research Foundation. C.P-L is supported by the Human Frontier Science Program Grant No. RGY0072/2021 and the University of Auckland Faculty of Science RDF #3732317. T.A.W. is supported by a grant from the Gordon and Betty Moore Foundation (GBMF9741) and the John Templeton Foundation (63451); the opinions expressed in this publication are those of the authors and do not necessarily reflect the views of the John Templeton Foundation.

## AUTHOR CONTRIBUTIONS

Conceptualization, B.M.C, M.D.H;

Methodology, M.L, F.B, K.E.C, E.J.S, R.J.C, B.M.C, M.D.H, E.R.R.M, T.A.W, C.P-L;

Investigation, M.L, F.B, K.E.C E.R.R.M, C.P-L, M.D.H, B.M.C;

Writing – Original Draft, B.M.C, M.D.H;

Writing – Review & Editing, M.L, F.B, K.E.C, E.J.S, R.J.C, B.M.C, M.D.H, E.R.R.M, T.A.W, C.P-L, K.A.M;

Funding Acquisition, R.J.C, B.M.C, M.D.H;

Supervision, B.M.C, M.D.H.

## CONFLICTS OF INTEREST

The authors declare no conflict of interest.

## Notes

### Competing Interest Statement

The authors have declared no competing interest.

### Summary of Updates

Changes have been made according to reviewer comments from Nature Communications.

