## Supplementary Data for "The prokaryotic origins of the COMMD protein family involved in eukaryotic membrane trafficking"

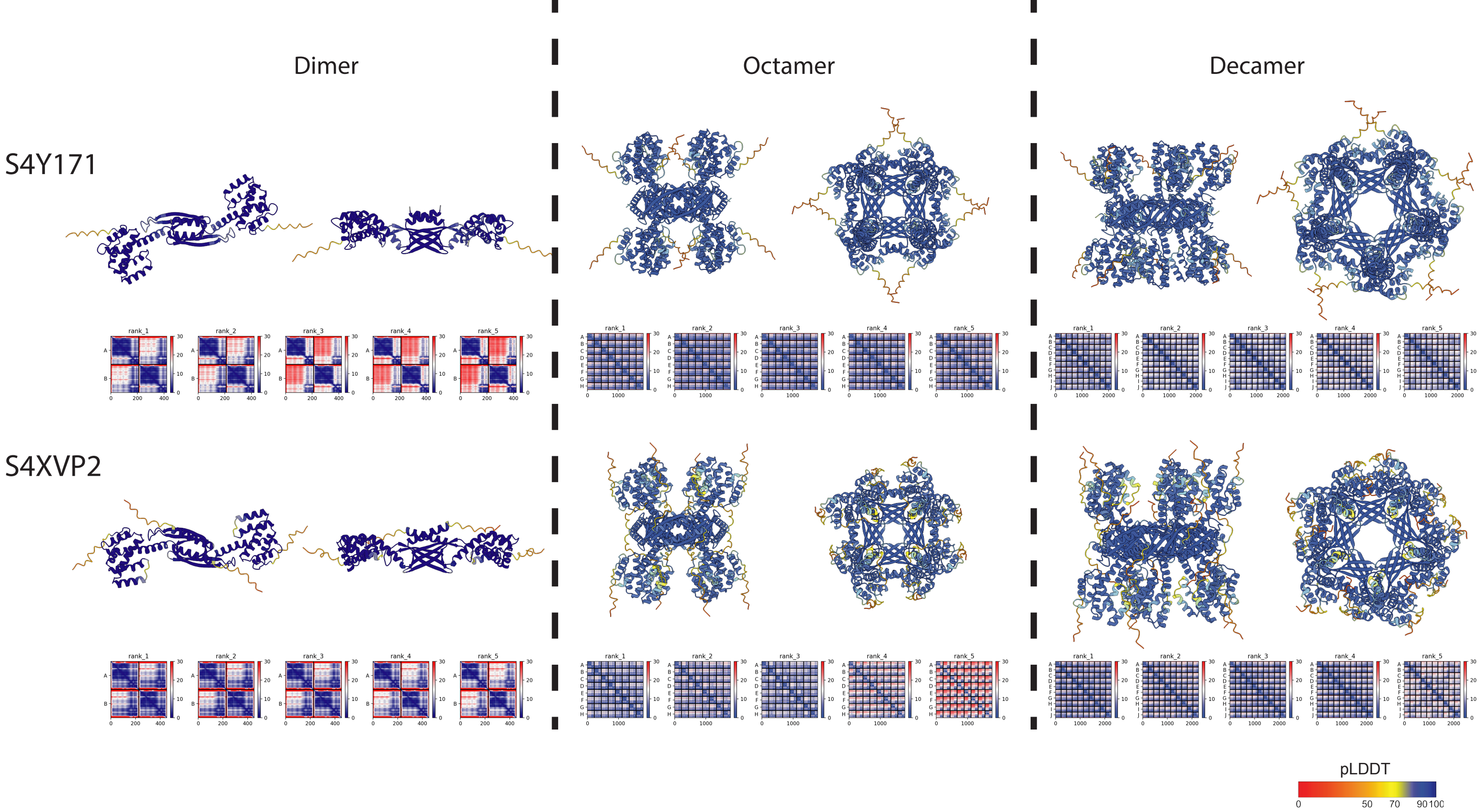

**Figure S1. AlphaFold2 predicted structures of COMMD-like proteins from Myxococcota.**
The structures of S4Y171 and S4XVP2 from Sorangium cellulosum were predicted using AlphaFold2 in various homooligomeric forms, with the results shown for dimer, octamer and decamer predictions. Proteins care coloured according to pLDDT, with PAE plots shown below.

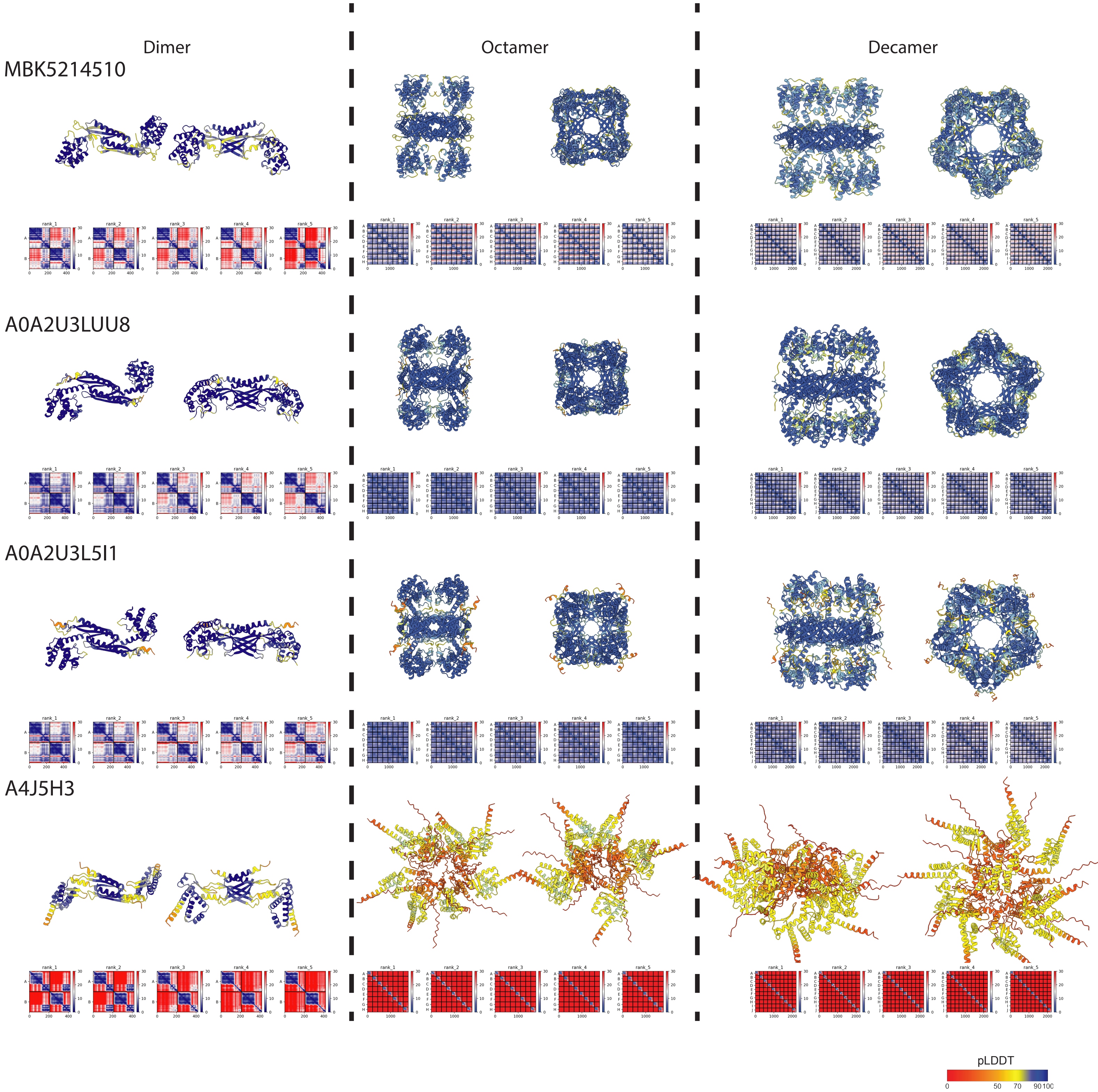

**Figure S2. AlphaFold2 predicted structures of COMMD-like proteins from various bacterial species.** The structures of A4J5H3 (Desulfotomaculum reducens), A0A2U3L5I1 (Candidatus Sulfopaludibacter), A0A2U3LUU8 (Candidatus Sulfopaludibacter), MBK5214510 (Flavobacteriaceae bacterium) were predicted using AlphaFold2 in various homooligomeric forms, with the results shown for dimer, octamer and decamer predictions. Proteins care coloured according to pLDDT, with PAE plots shown below.

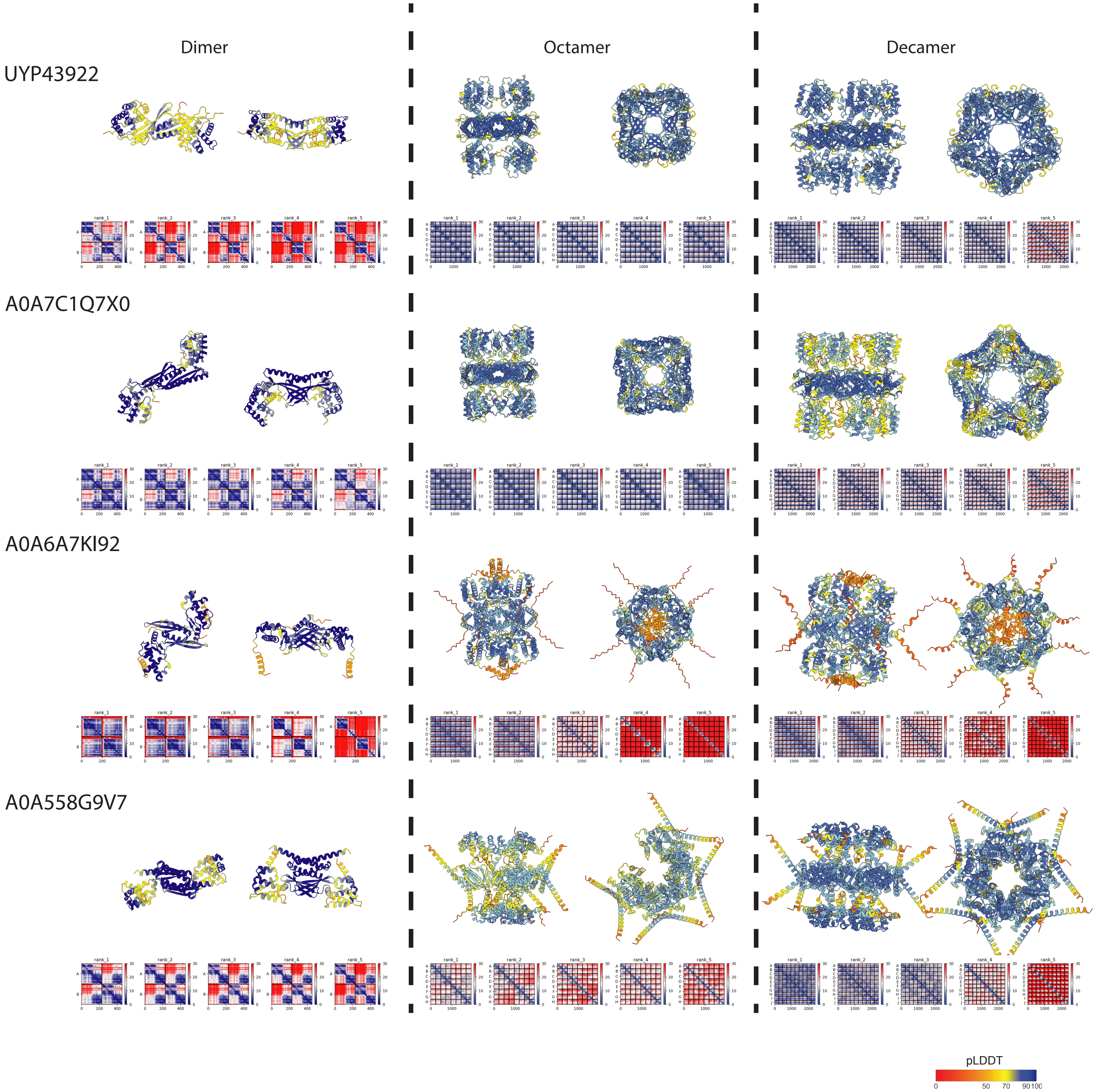

**Figure S3. AlphaFold2 predicted structures of COMMD-like proteins from various archaeal species.** The structures of UYP43922 (Candidatus Lokiarchaeum ossiferum), A0A7C1Q7X0 (Unclassified), A0A6A7KI92 (Thermococcus sp. 101 C5), A0A558G9V7 (Haloferax volcanii) were predicted using AlphaFold2 in various homooligomeric forms, with the results shown for dimer, octamer and decamer predictions. Proteins care coloured according to pLDDT, with PAE plots shown below.

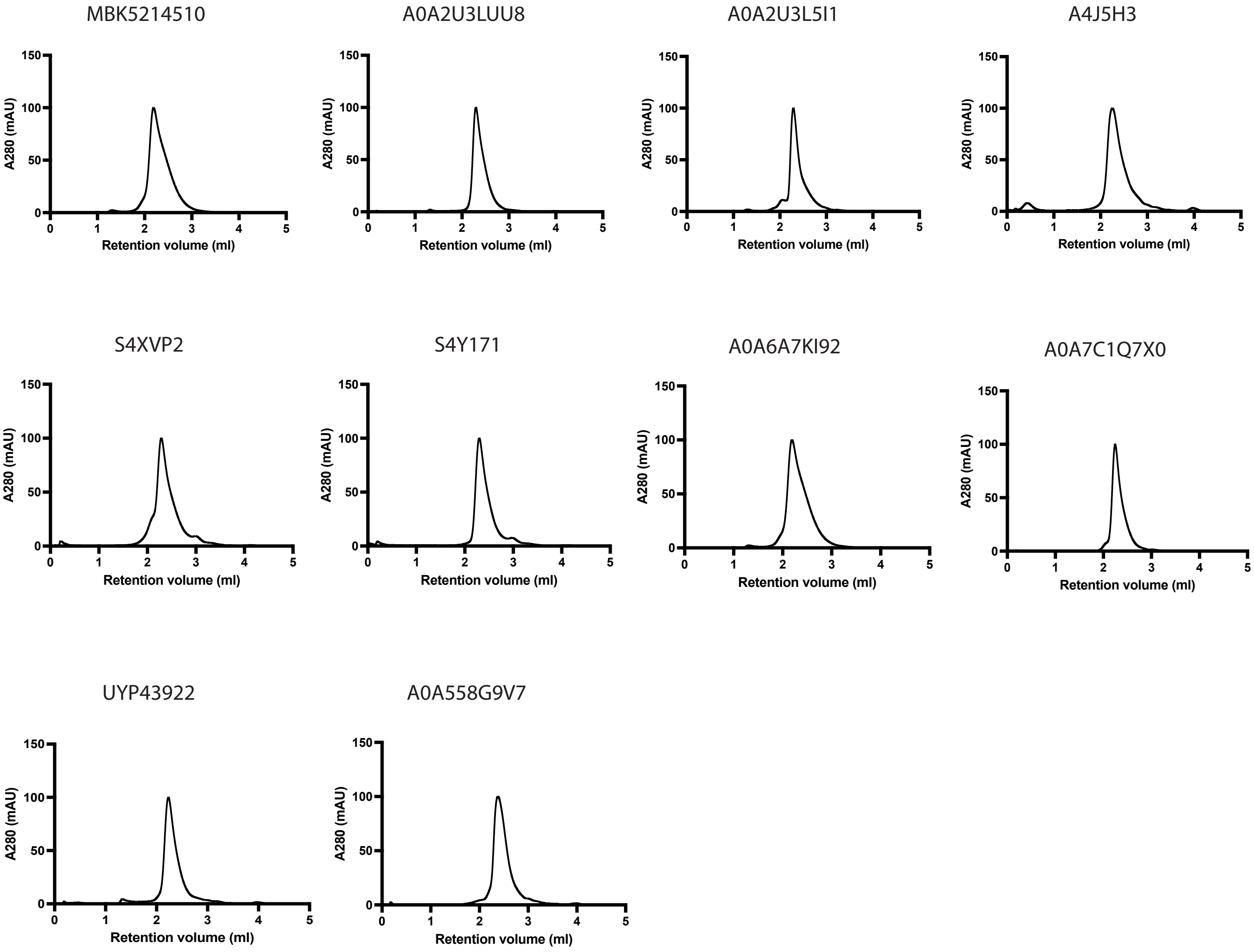

**Figure S4. Size exclusion chromatography profiles of COMMD-like proteins.**
All proteins were purified on a superdex 200 10/150 column attached to an AKTA pure system. This yielded proteins with relatively homogenous peaks.

**A**

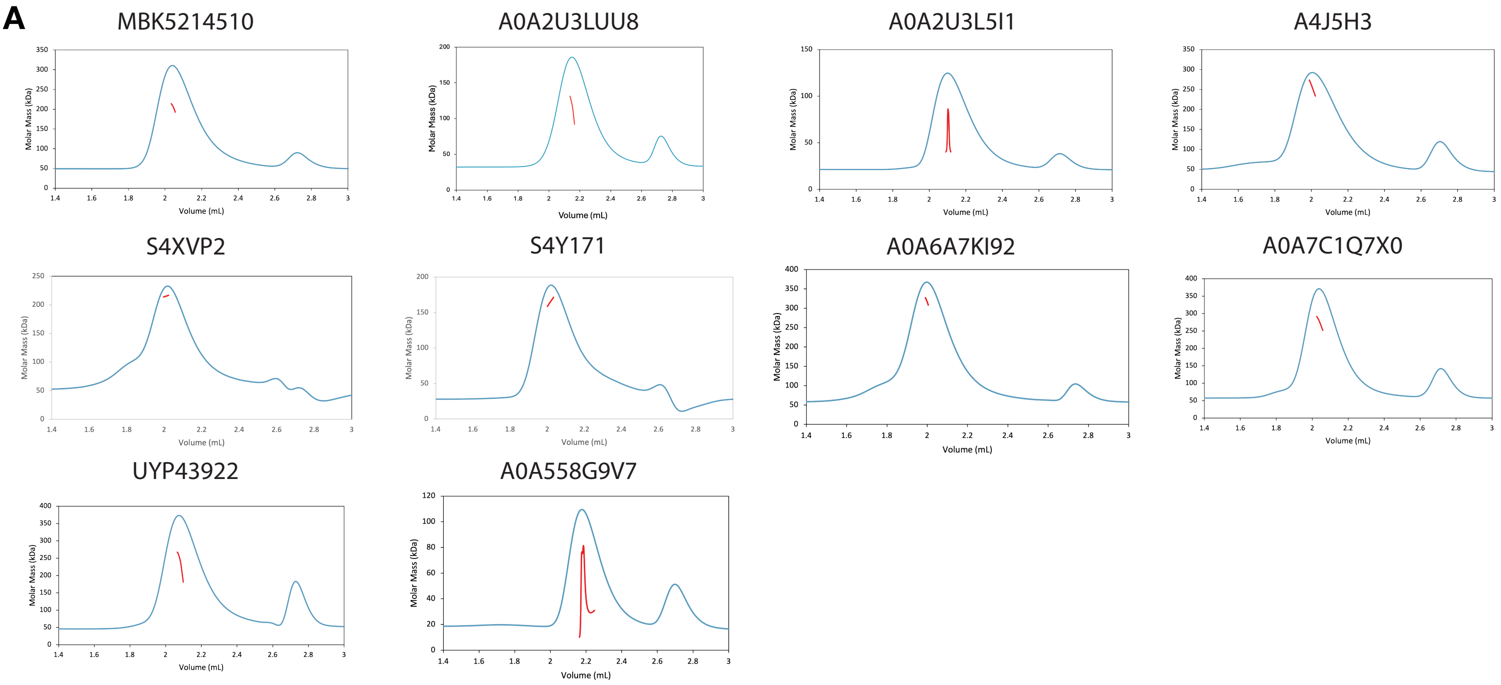

**B**

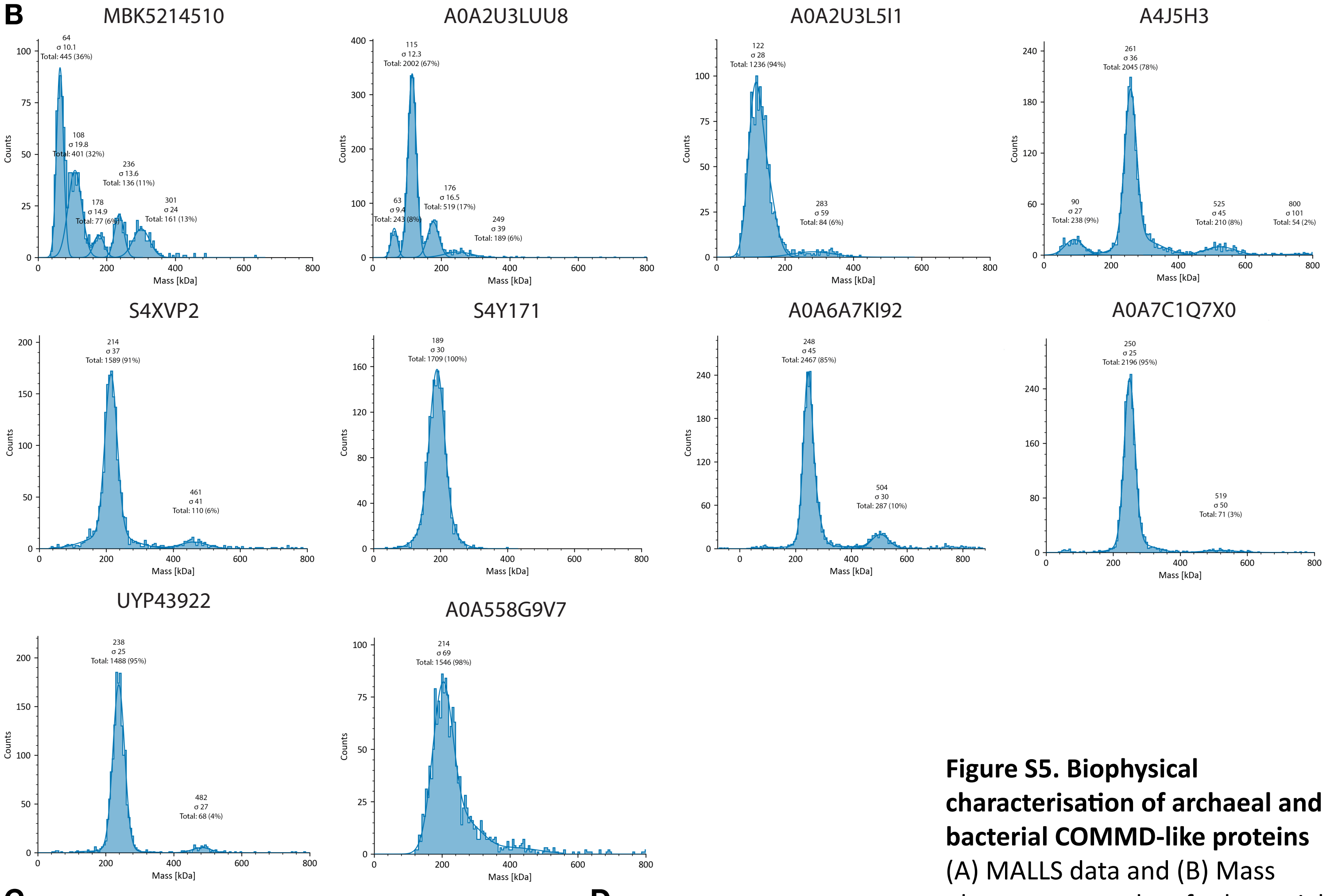

**C**

| Kingdom | Uniprot | Monomer MW (kDa) |
|---|---|---|
| Bacteria | MBK5214510 | 27.7 |
| | A0A2U3LUU8 | 28.2 |
| | A0A2U3L5I1 | 29.6 |
| | A4J5H3 | 25.3 |
| | S4XVP2 | 25.5 |
| | S4Y171 | 23.6 |
| Archaea | A0A6A7KI92 | 24.0 |
| | A0A7C1Q7X0 | 29.5 |
| | UYP43922 | 28.5 |
| | A0A558G9V7 | 27.8 |

**D**

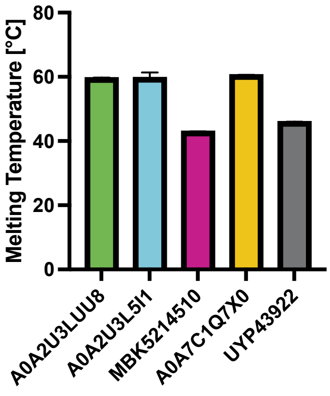

**Figure S5. Biophysical characterisation of archaeal and bacterial COMMD-like proteins** (A) MALLS data and (B) Mass photomometry data for bacterial and archaeal COMMD-like proteins. (C) expected molecular weight of COMMD-like proteins including purification tags and cloning scars. (D) NanoDSF shows two groupings of COMMD-like proteins with respect to thermal stability with a TM50 of ~60°C and 40°C.

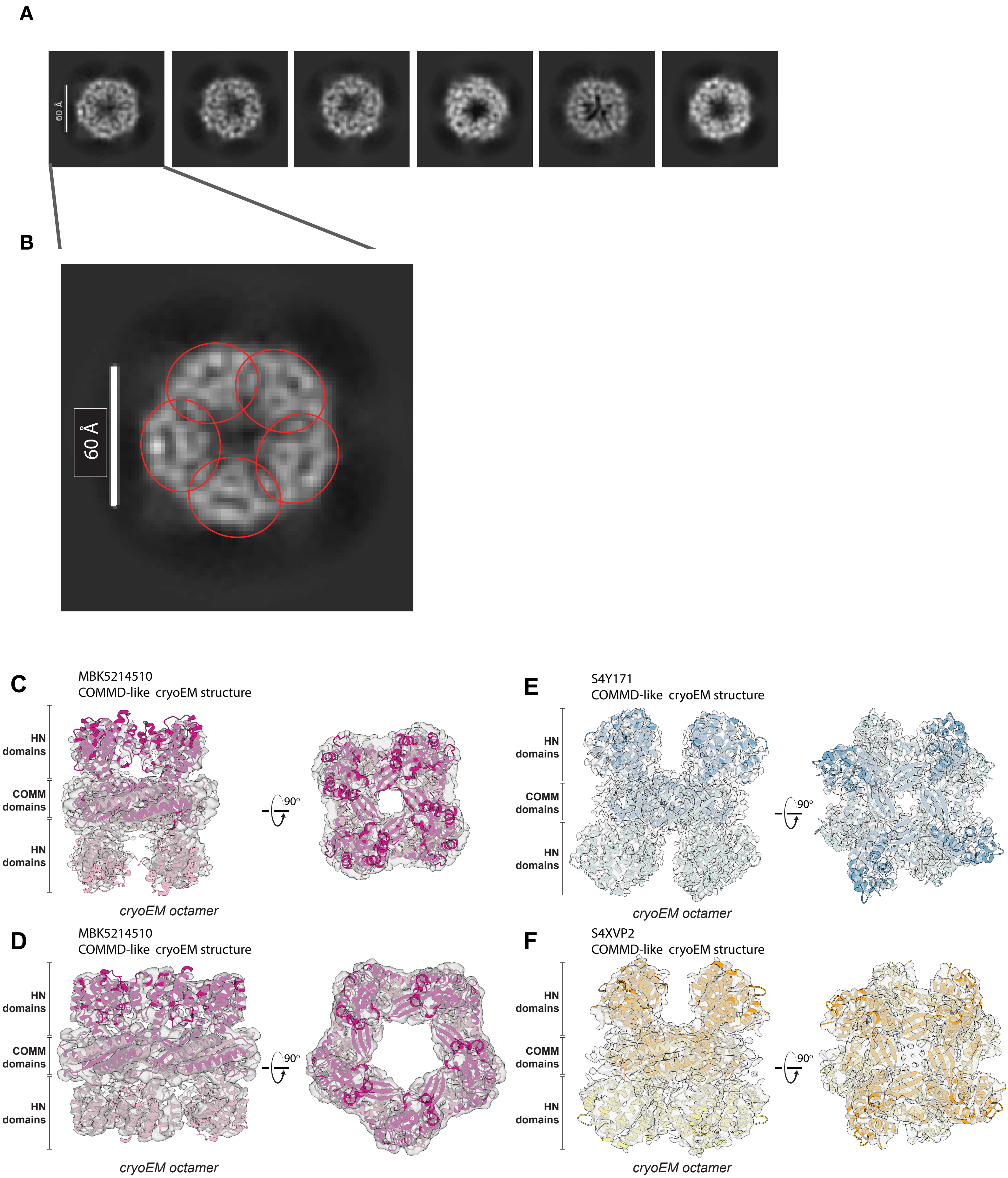

**Figure S6. Cryo-EM of COMMD-like proteins.**
(A) 2D class averages indicating a decameric complex of COMMD-like protein A0A6A7KI92 from Thermococcus sp. 101 C5. (B) Highlight of the 5 protein dimers in the 2D classified top view. (C-F) Density maps with fitted structure of the MBK5214510 octamer, the MBK5214510 decamer, the S4Y171 octamer and the S4XVP2 octamer respectively.

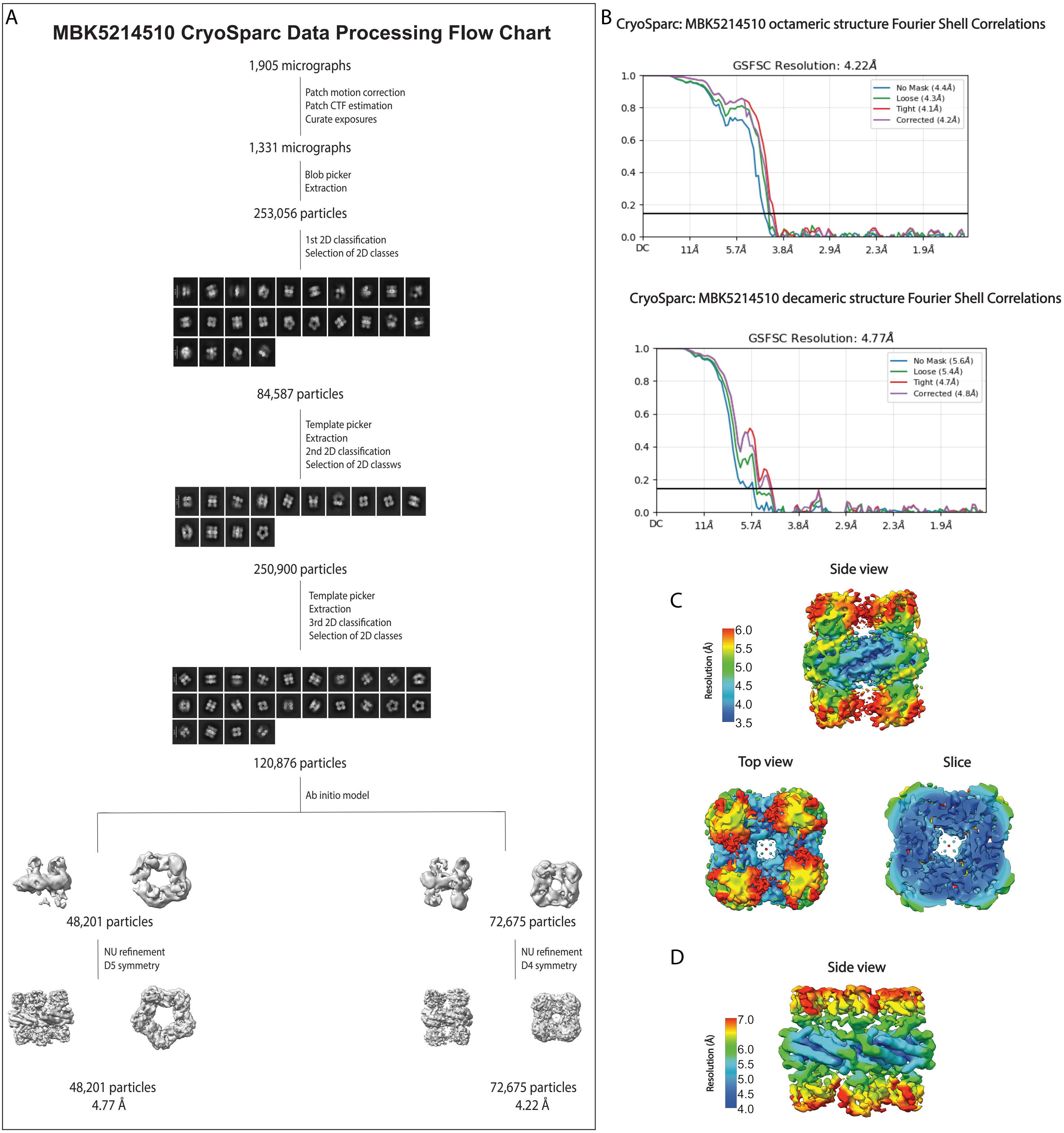

**Figure S7. Cryo-EM workflow for structure determination of MBK5214510.**

(A) Cryo-EM data processing workflow using CryoSPARC software. The final 3D reconstruction map was used for model building and refinement. (B) 3D Fourier Shell Correlation of the final reconstructions reporting global resolution, with the FSC at 0.143 denoted. (C) Final reconstruction of octameric structure filtered and colored by local resolution, as estimated in CryoSPARC. (D) Final reconstruction of decameric structure filtered and colored by local resolution, as estimated in CryoSPARC.

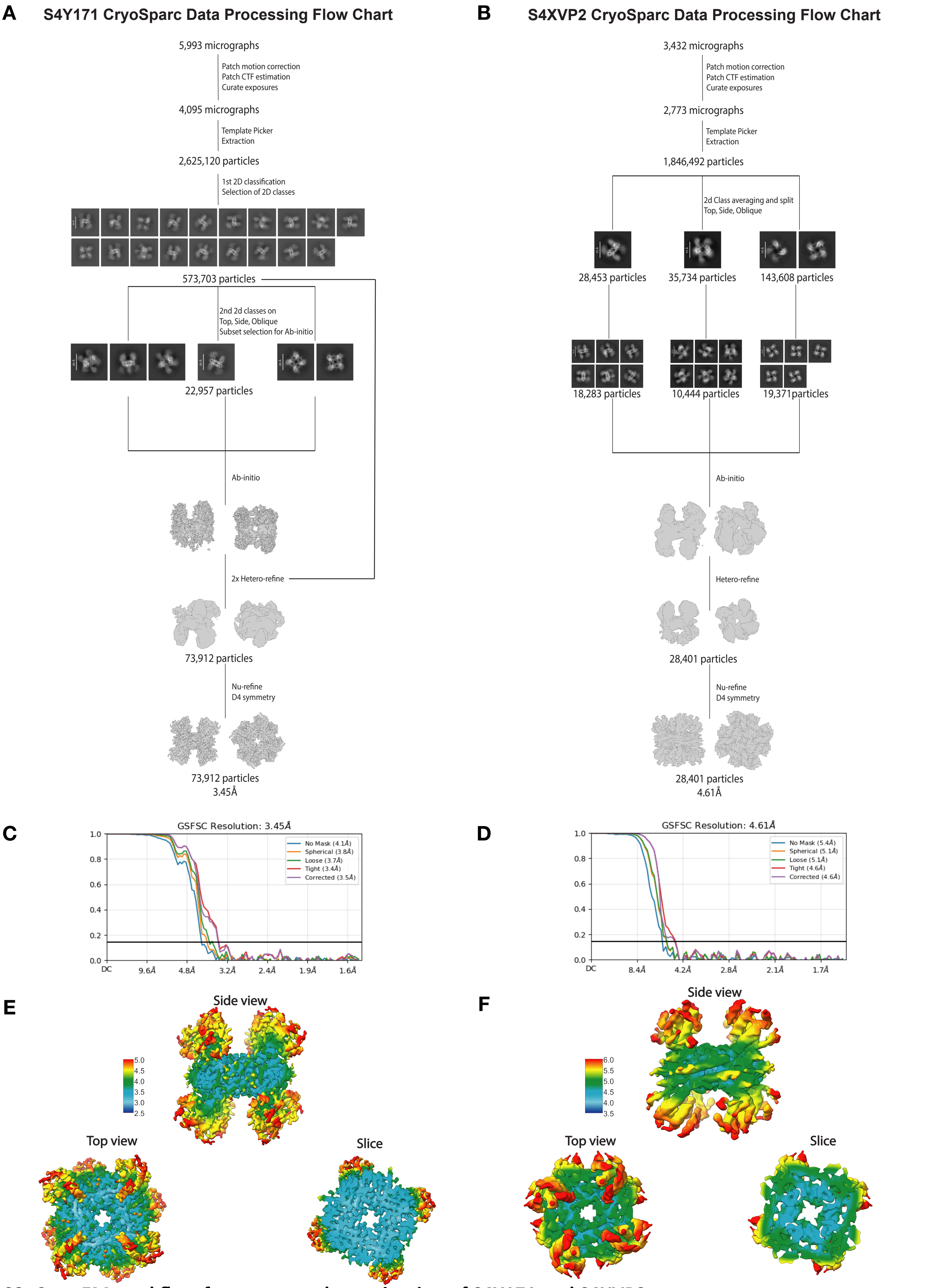

**Figure S8. Cryo-EM workflow for structure determination of S4Y171 and S4XVP2.**
(A) and (B) Cryo-EM data processing workflow using CryoSPARC software. The final 3D reconstruction map was used for model building and refinement. (C) and (D) 3D Fourier Shell Correlation of the final reconstructions reporting global resolution, with the FSC at 0.143 denoted. (E) Final reconstruction of the S4Y171 octameric structure filtered and colored by local resolution, as estimated in CryoSPARC. (F) Final reconstruction of the S4XVP2 octameric structure filtered and colored by local resolution, as estimated in CryoSPARC.

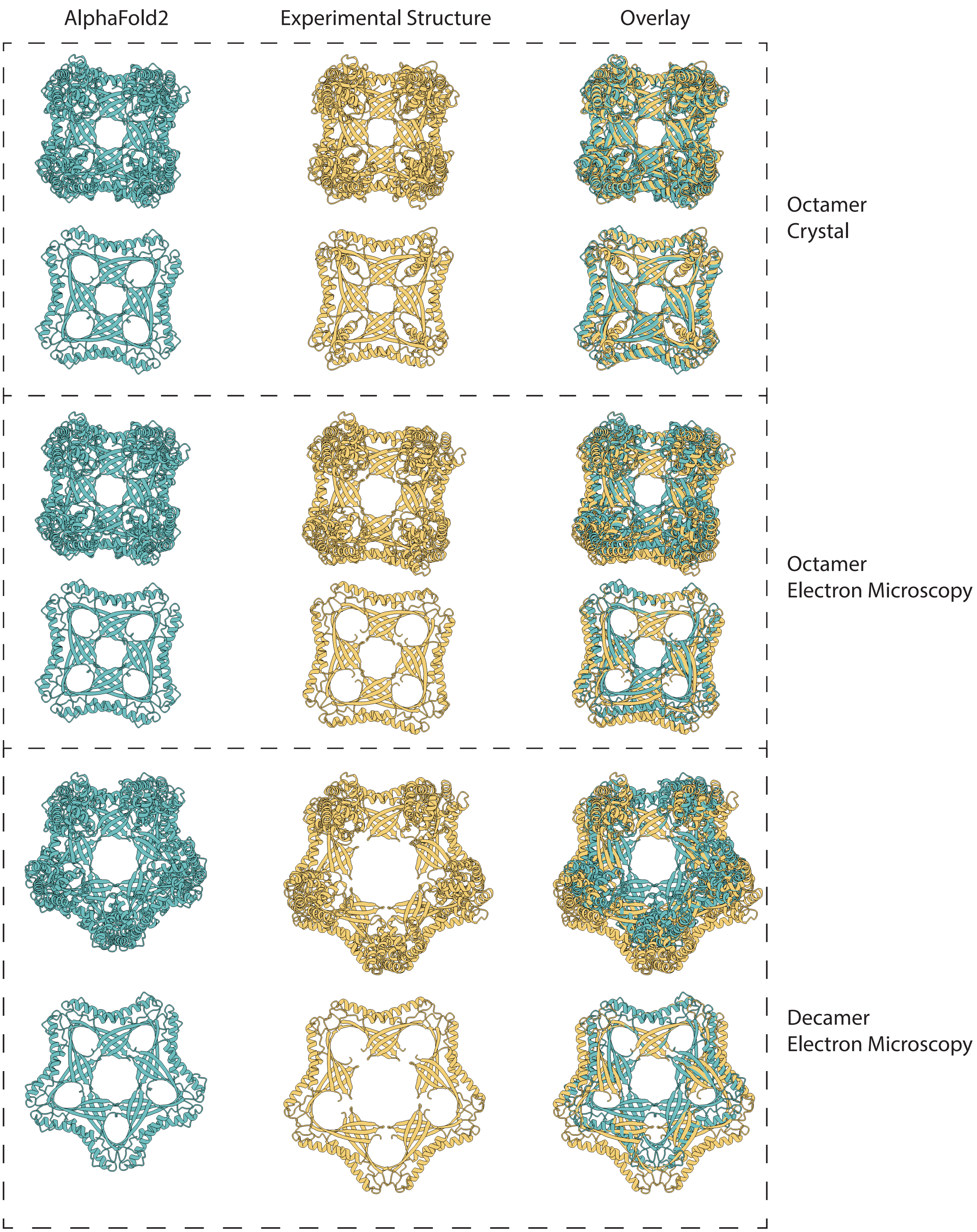

**Figure S9. Comparison of the experimental MBK5214510 COMMD-like structures with the AlphaFold predictions.**
Left panels show AlphaFold2-predicted structures of MBK5214510 COMMD-like protein from Flavobacteriaceae bacterium with the lower images showing just the central ring of COMM domains that form the core of the homooligomeric rings. Middle panels show the experimental structures of MBK5214510 octamers and decamers determined by X-ray crystallography and cryo-EM and right-hand panels show the overlay of the predicted and experimental structures.

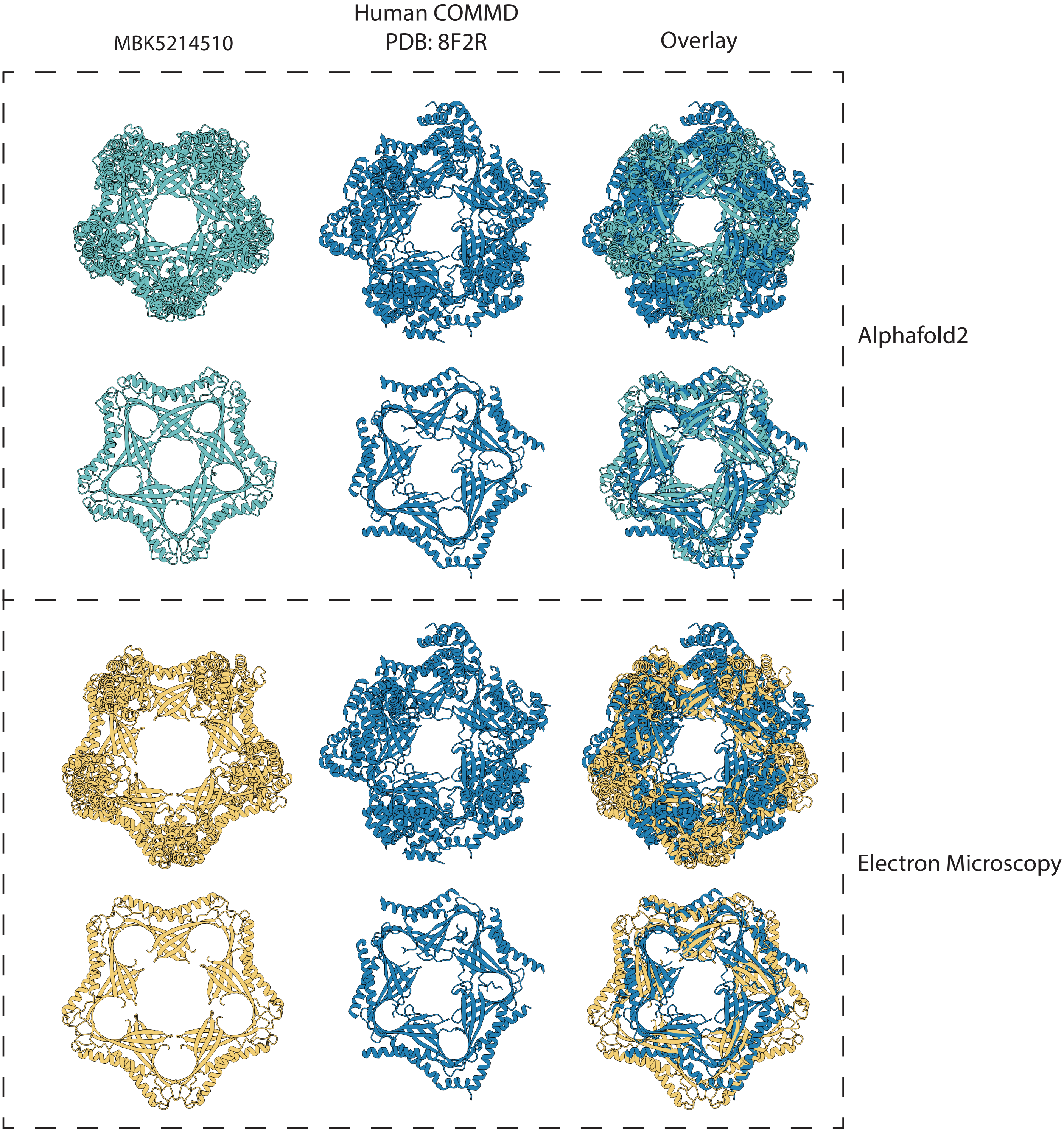

**Figure S10. Comparison of the experimental MBK5214510 COMMD-like homodecameric structure with the human COMMD heterodecameric complex.**
Left panels show either the AlphaFold2-predicted or cryo-EM homodecameric structure of MBK5214510 COMMD-like protein from Flavobacteriaceae bacterium with the lower images showing just the central ring of COMM domains that form the core of the homooligomeric rings. Middle panels show either the AlphaFold2-predicted or cryo-EM heterodecameric structure the human COMMD1-COMMD10 complex18 (PDB ID 8F2U) and right-hand panels show the overlay of the bacterial and human structures.

**A**

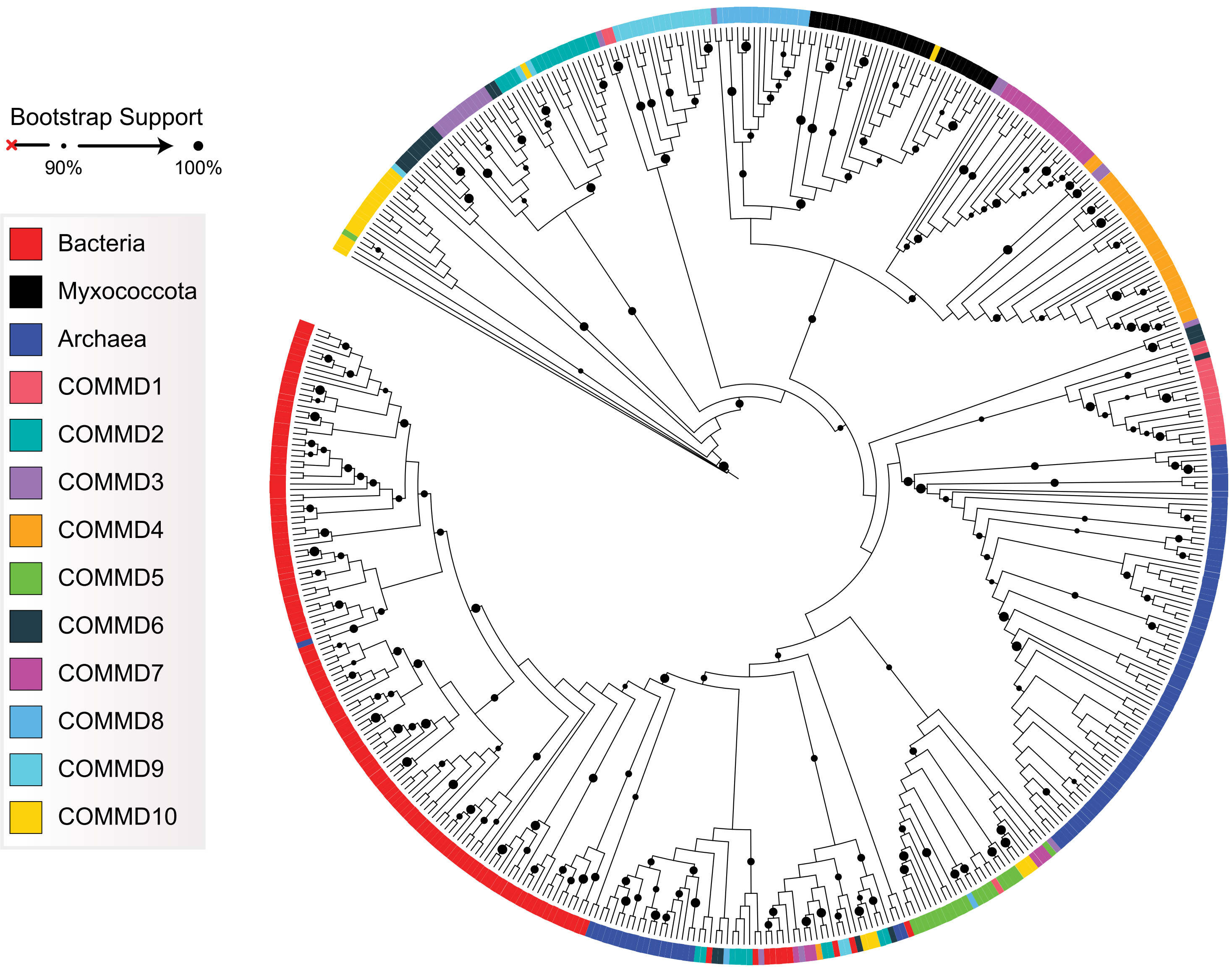

**B**

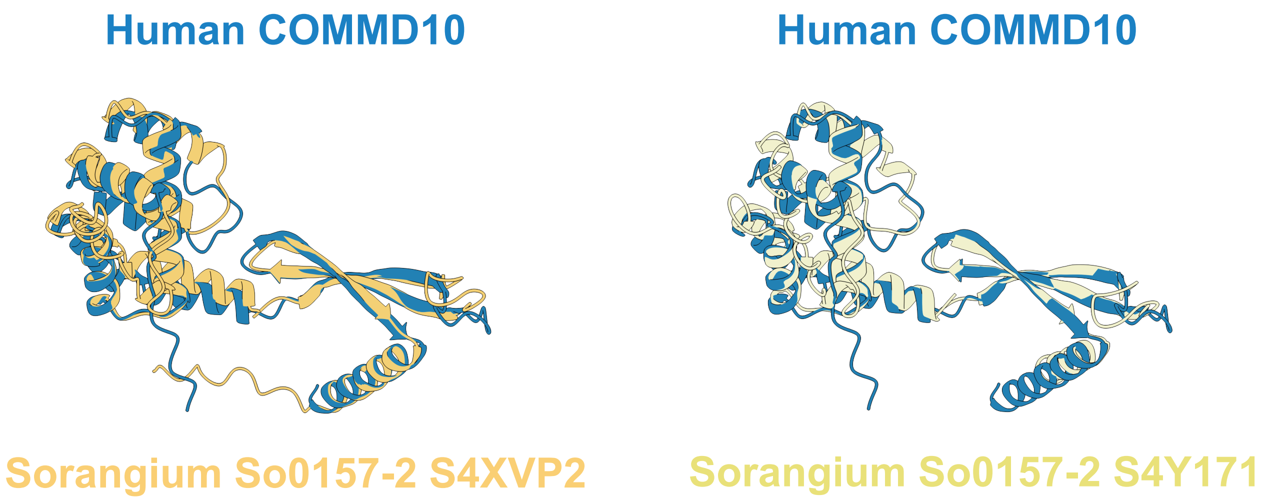

**Figure S11. Structure-based phylogenetics.**

(A) 3Di phylogeny showing the relationship between archaeal (blue), bacterial (red) Myxococcota (light green) COMMD-like proteins and eukaryotic COMMD1-COMMD10 proteins. Bootstraps support between 90% and 100% is noted by a circle of increasing size. Branches with bootstrap values below 90% are considered poorly support and therefore not indicated. (B) Example structural alignment of the AlphaFold2 predictions of COMMD10 with two Mycococcota COMMD-like proteins S4Y171 and S4XVP2 from Sorangium cellulosum.

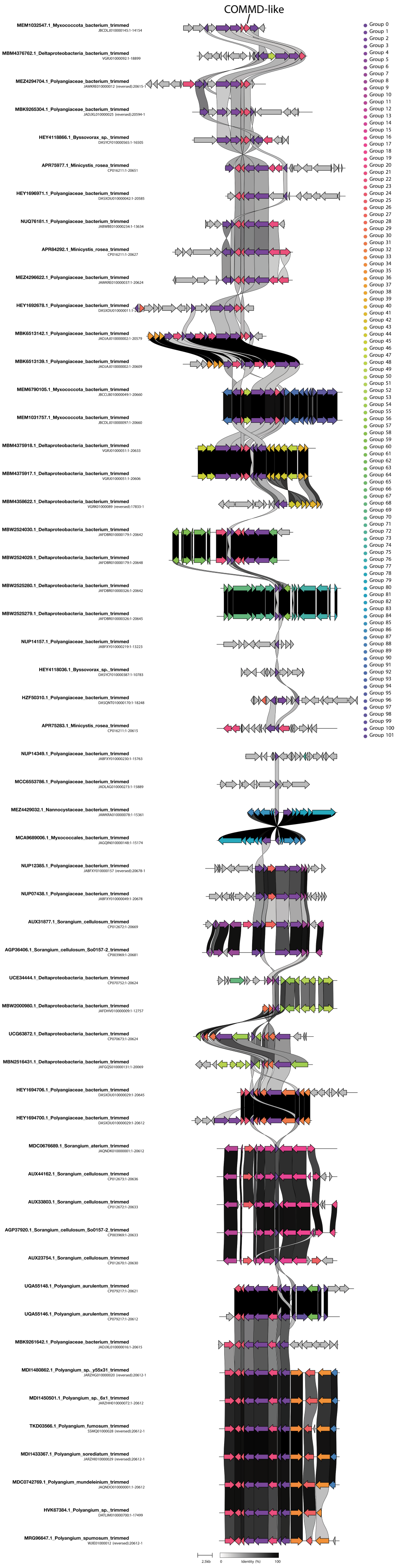

**Figure S12. Gene clusters surrounding COMMD-like genes in Myxococotta.**
Analysis of the genomic surroundings of COMMD-like genes from Myxococotta species using CLINKER.

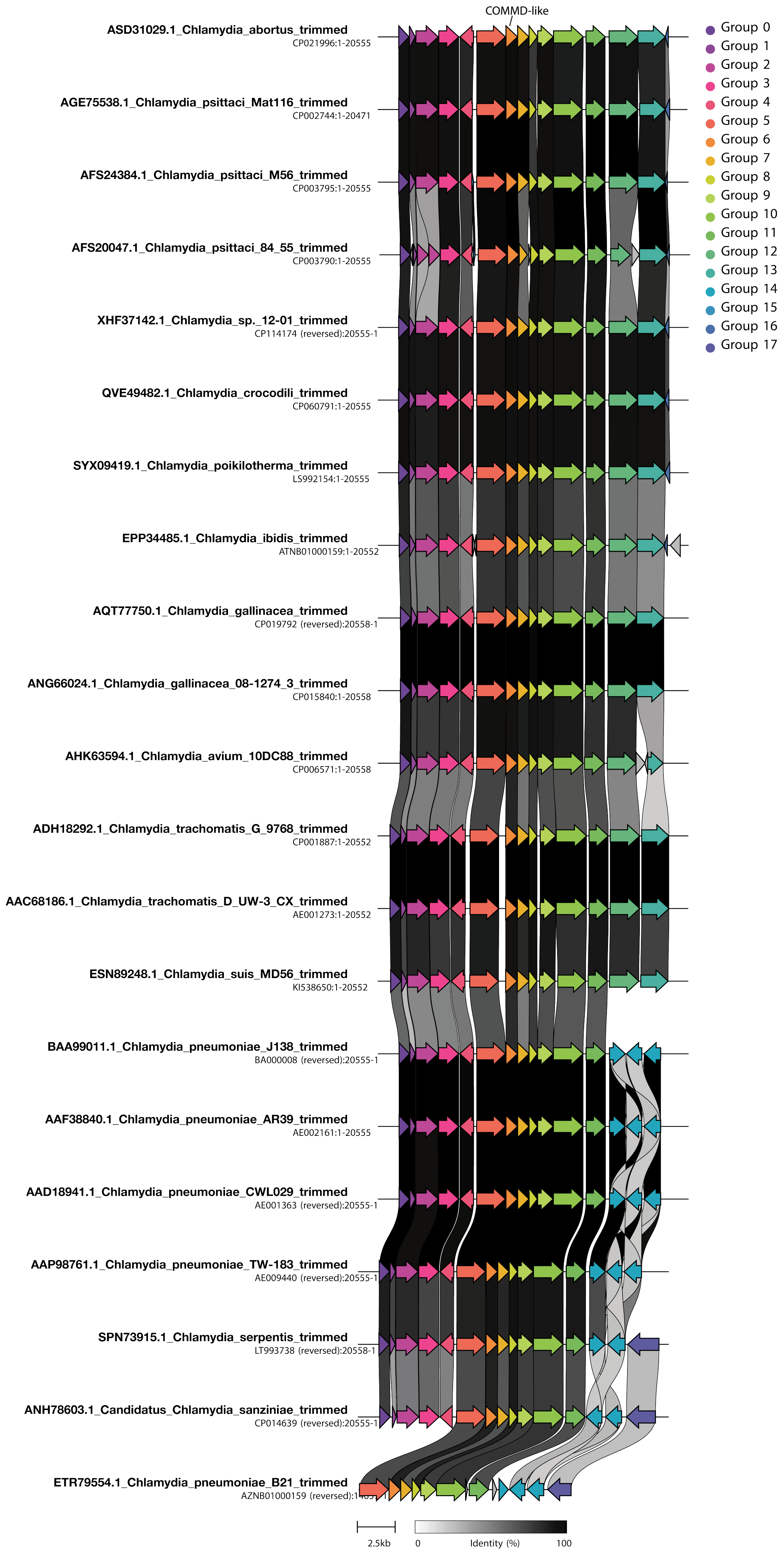

**Figure S13. Gene clusters surrounding COMMD-like genes in Chlamydia.**
Analysis of the genomic surroundings of COMMD-like genes from Chlamydial species using CLINKER.

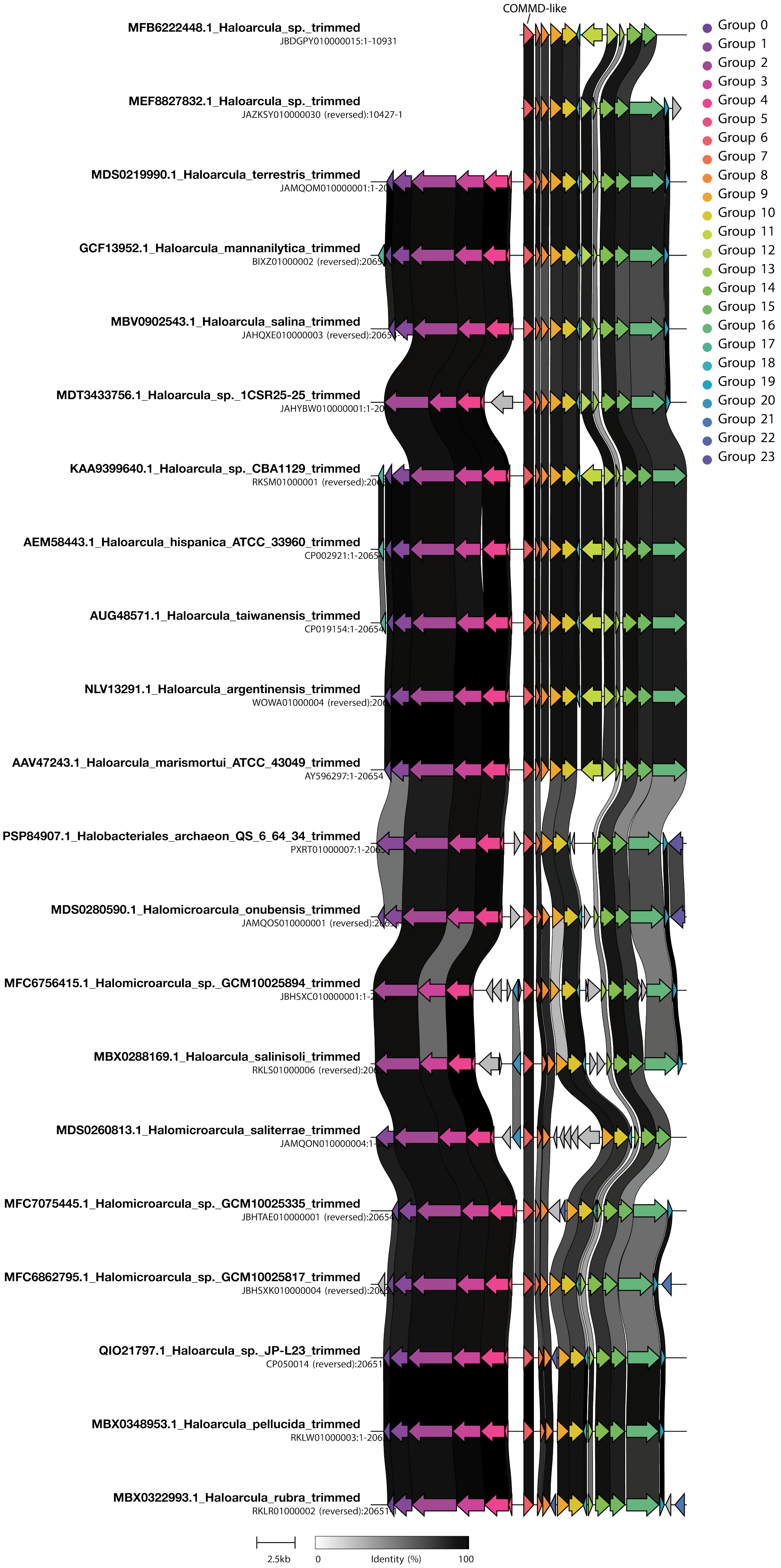

**Figure S14. Gene clusters surrounding COMMD-like genes in Haloarchaea.**
Analysis of the genomic surroundings of COMMD-like genes from Haloarchaea species using CLINKER.

**Supplementary Table S1.** Crystallographic data collection and refinement statistics.

| | **MBK5214510** |
|---|---|
| **Data Collection** | |
| Space group | I222 |
| Unit cell dimensions (a,b,c) (Å) | 91.3, 96.0, 115.6 |
| Unit cell angles (α,β,γ) (º) | 90 90 90 |
| Wavelength (Å) | 0.95373 |
| Resolution range (Å) | 45.67 - 2.582 (2.66 - 2.58) |
| Total reflections | 106202 |
| Unique reflections | 16103 (1127) |
| Multiplicity | 6.6 (6.4) |
| Completeness (%) | 98.69 (84.74) |
| Mean I/σ(I) | 14.8 (2.8) |
| R-merge | 0.022 (0.146) |
| R-meas | 0.0313 (0.207) |
| R-pim | 0.022 (0.146) |
| CC(1/2) | 0.999 (0.919) |
| **Refinement** | |
| R-work | 0.2419 (0.2919) |
| R-free | 0.2895 (0.3782) |
| Number of non-hydrogen atoms | 3573 |
| Number macromolecule atoms | 3534 |
| Number water atoms | 39 |
| RMS bonds (Å) | 0.006 |
| RMS angles (º) | 1.17 |
| Ramachandran favored (%) | 93.44 |
| Ramachandran allowed (%) | 4.92 |
| Ramachandran outliers (%) | 1.64 |
| Clashscore | 10.97 |
| Overall Molprobity score | |
| Average B-factor | 71.52 |
| PDB ID | 9Q6A |

**a.** Statistics for the highest-resolution shell are shown in parentheses.

**Supplementary Table S2.** Cryo-EM data collection and refinement statistics.

| | **MBK5214510 octamer** | **MBK5214510 decamer** | **S4Y171** | **S4XVP2** |
|---|---|---|---|---|
| **Data collection and processing** | | | | |
| Magnification | 100,000 | 100,000 | 190,000 | 190,000 |
| Voltage (kV) | 300 | 300 | 200 | 200 |
| Electron exposure ($e^{-}/Å^2$) Defocus range (µm) | 40 $e^{-}/Å^2$ -2.5 µm | 40 $e^{-}/Å^2$ -2.5 µm | 50 $e^{-}/Å^2$ -2.5 µm | 50 $e^{-}/Å^2$ -2.5 µm |
| Pixel size (Å) | 0.82 | 0.82 | 0.75 | 0.75 |
| Symmetry imposed | D4 | D5 | D4 | D4 |
| Programme used | CryoSPARC | CryoSPARC | CryoSPARC | CryoSPARC |
| Initial micrographs collected (no.) | 1,905 | 1,905 | 5,993 | 3,432 |
| Final micrographs processed (no.) | 1,331 | 1,331 | 610 | 2,773 |
| Initial particle images (no.) | 253,056 | 253,056 | 2,625,120 | 1,846,492 |
| Final particle images (no.) | 72,675 | 48,201 | 73,912 | 28,401 |
| Map resolution (Å) | 4.22 | 4.77 | 3.45 | 4.61 |
| FSC threshold | 0.143 | 0.143 | 0.143 | 0.143 |
| **Refinement** | | | | |
| Initial model used | ColabFold | ColabFold | ColabFold | ColabFold |
| Model resolution range (0/0.143/0.5) (Å) | 4.2/4.3/7.7 | 4.6/4.7/7.6 | 3.1/3.4/4.2 | 4.6/4.7/5.3 |
| **Model composition** | | | | |
| Non-hydrogen atoms | 14,368 | 17,960 | 24,374 | 24,568 |
| Protein residues | 1760 | 2200 | 1584 | 1584 |
| R.m.s. deviations | | | | |
| Bond lengths (Å) | 0.008 | 0.002 | 0.003 | 0.004 |
| Bond angles (°) | 0.861 | 0.548 | 0.555 | 0.513 |
| Validation | | | | |
| MolProbity score | 2.17 | 1.86 | 1.45 | 1.64 |
| Clashscore | 19.74 | 12.81 | 4.39 | 13.49 |
| **Ramachandran plot** | | | | |
| Favoured (%) | 94.5 | 96.33 | 96.49 | 98.47 |
| Allowed (%) | 5.5 | 3.67 | 3.51 | 1.53 |
| Outliers (%) | 0 | 0 | 0 | 0 |
| Database Information | | | | |
| EMDB ID | EMD-72266 | EMD-72267 | EMD-73963 | EMD-73964 |
| PDB ID | 9Q6C | 9Q6D | 9ZA1 | 9ZA2 |

**Supplementary Table S3.** Key Resources Table.

| REAGENT or RESOURCE | SOURCE | IDENTIFIER |
|---|---|---|
| **Bacterial strains** | | |
| *E. coli* DH5α | Invitrogen | Cat# 18265017 |
| BL21 (DE3) | New England Biolabs | Cat# C2527H |
| DH10EMBacY | ThermoFisher | Cat# 10361012 |
| NEB 5-alpha | New England Biolabs | Cat# C2987H |
| **Chemicals and peptides** | | |
| Benzamidine hydrochloride hydrate | Sigma Aldrich | Cat# B6506 |
| Deoxyribonuclease I (DNase I) | Sigma Aldrich | Cat# DN25 |
| Talon® resin | Clontech | Cat# 635503 |
| Glutathione Sepharose 4B | GE Healthcare | Cat# GEHE17-0756-0 |
| Isopropyl β-D-1-thiogalactopyranoside | Bioline | Cat# BIO-37036 |
| Triton-X100 | Sigma Aldrich | Cat# X100-500ML |
| Sf-900 II SFM media | ThermoFisher Scientific | Cat# 10902088 |
| Graphene oxide | Sigma-Aldrich | Cat# 763705 |
| **Critical commercial assays** | | |
| QIAprep Spin Miniprep Kit | QIAGEN | Cat# 27106 |
| NEBuilder HiFi DNA Assembly System (NEB) | New England Biolabs | Cat# E5520S |
| TMT seven-plex | ThermoFisher | Cat# 90061 |
| **Recombinant DNA** | | |
| pET-28(a) | Sigma-Aldrich | Cat# 69864 |
| LM627 vector | Addgene | 191661 |
| **Deposited Data** | | |
| MBK5214510 (crystal structure) | RCSB Protein Data Bank (this study) | PDB ID: 9Q6A |
| MBK5214510 octamer (cryo-EM structure; RELION map) | RCSB Protein Data Bank (this study) | PDB ID: 9Q6C<br>EMDB ID: EMD-72266 |
| MBK5214510 decamer (cryo-EM structure; CryoSPARC map) | RCSB Protein Data Bank (this study) | PDB ID: 9Q6D<br>EMDB ID: EMD-72267 |
| S4Y171 Octamer (cryo-EM structure; CryoSPARC map) | RCSB Protein Data Bank (this study) | PDB ID: 9ZA1<br>EMDB ID: EMD-73963 |
| S4XVP2 Octamer (cryo-EM structure; CryoSPARC map) | RCSB Protein Data Bank (this study) | PDB ID: 9ZA2<br>EMDB ID: EMD-73964 |
| AlphaFold2 models of Bacterial and Archaeal COMMD-like proteins | ModelArchive (this study) | 10.5452/ma-commd-like |
| **Software and Algorithms** | | |
| XDS | [1] | http://xds.mpimf-heidelberg.mpg.de/ |
| AIMLESS | [2] | http://www.ccp4.ac.uk/html/aimless.html |
| Phaser | [3] | http://www.phaser.cimr.cam.ac.uk/index.php/Phaser_Crystallographic_Software |

| | | |
|---|---|---|
| Phenix | [4] | https://www.phenix-online.org/ |
| Coot | [5] | https://www2.mrc-lmb.cam.ac.uk/personal/pemsley/coot/ |
| ChimeraX | [6] | https://www.rbvi.ucsf.edu/chimerax/ |
| iSOLDE | [7] | https://isolde.cimr.cam.ac.uk |
| CryoSPARC V4.7.1 | [8] | https://cryosparc.com |
| AlphaFold2 Multimer | [9] | https://github.com/deepmind/alphafold |
| AlphaFold3 | | |
| ColabFold v1.5.5 implementation of AlphaFold2-Multimer | [10] | https://github.com/sokrypton/ColabFold |
| FoldSeek version c27a629a0a345c288c18d2a0975a3dc7bb143798 | [11] | https://github.com/steinegglab/foldseek |
| MAFFT v7.505 | [12] | https://mafft.cbrc.jp/alignment/server/index.html |
| IQTREE 2.2.6 | [13] | https://iqtree.github.io |
| FOLDMASON | [14] | https://github.com/steinegglab/foldmason |
| Figtree v1.4.4 | | http://tree.bio.ed.ac.uk/software/figtree/ |
| CLINKER | [15] | https://github.com/gamcil/clinker |
| **Other** | | |
| Superose6 Increase10/300 GL column | Cytiva | Cat# 29091596 |
| HiLoad™ Superdex75 PG column | Cytiva | Cat# 28989333 |
| MonoQ 5/50 GL column | Cytiva | Cat# 17516601 |
| C-Flat 1.2/1.3 grids | C-Flat | N/A |
| Quantifoil 1.2/1.3 Copper 300 mesh grids | Quantifoil | N/A |

**Supplementary Table S4.** *Myxococcota* genomic clustering

| CDS | Closest PDB | Description |
|---|---|---|
| APR75977.1 | 8F2U[16] | Human Decameric COMMD ring |
| MEZ4294706.1 | | |
| MEZ4294709.1 | | |
| MBM4375919.1 | 6H3N[17] | VgrG1 of *Pseudomonas aeruginosa PA01* |
| APR75972.1 | 6RBK[18] | anti-feeding prophage (AFP) baseplate |
| APR84297.1 | | |
| HEY1692683.1 | 4JIW[19]/7RPQ[20] | PAAR(N-terminus)/ctpa protease (C-term) |
| HEY1696975.1 | 4EQM[21] | Serine-Threonine kinase |
| NUP07431.1 | 6KON[22]/2PL2[23] | Component of *Mycobacterium tuberculosis* initial transcription complex/ tetratricopeptide repeat |
| MBM4375922.1 | 2M2K[24] | HasB |
| UCE34441.1 | 3KRU[25] | Old Yellow Enzyme |

**Supplementary Table S5.** *Chlaymidia* and *Haloarchaea* example protein clusters

| CDS | Uniprot | Closest PDB | Description |
|---|---|---|---|
| Chlamydia trachomatis serovar D (strain ATCC VR-885 / DSM 19411 / UW-3/Cx) | | | |
| AAC68178.1 | O84580 | 2VGX[26] | T3S Chaperon |
| AAC68179.1 | O84581 | | |
| AAC68180.1 | O84582 | | |
| AAC68181.1 | O84583 | | |
| AAC68182.1 | O84584 | 5I20[27] | DMT transporter |
| AAC68183.2 | O84585 | 1QF6[28] | THREONYL-TRNA SYNTHETASE |
| AAC68184.1 | O84586 | 6IUD[29] | DNA-partition |
| AAC68185.1 | O84587 | | |
| AAC68186.1 | O84588 | 3Q9D | COMMD-Like |
| AAC68187.1 | O84589 | 2A4M[30] | Tyrptophan synthase |
| AAC68188.1 | O84590 | 6O8E[31] | DNA repair/URVB |
| AAC68189.1 | O84591 | 3ZLF[32] | Enolase |
| AAC68190.1 | O84592 | 6MAB[33] | Periplasmic environment sensor |
| AAC68191.1 | O84593 | 3W43[34]/5LTV[35] | Stress signaling/receptor |
| Haloarcula hispanica (strain ATCC 33960 / DSM 4426 / JCM 8911 / NBRC 102182 / NCIMB 2187 / VKM B-1755) | | | |
| AEM58436.1 | G0HSR0 | | |
| AEM58437.1 | G0HSR1 | 2CY1[36] | Transcription termination factor |
| AEM58438.1 | G0HSR2 | 7OQ4[37] | RPO1C |
| AEM58439.1 | G0HSR3 | 7OQ4[37] | RPO1N |
| AEM58440.1 | G0HSR4 | 7OQ4[37] | RPO2 |
| AEM58441.1 | G0HSR5 | 7OQ4[37] | RPO2 |
| AEM58442.1 | G0HSR6 | 7OQ4[37] | RPO5 |
| AEM58443.1 | G0HSR7 | 3Q9D | COMMD-LIKE |
| AEM58444.1 | G0HSR8 | 7OYK[38] | Transcription regulator |
| AEM58445.1 | G0HSR9 | 6XS5[39] | VPS29 |
| AEM58446.1 | G0HSS0 | 7ULE[40] | F420-gamma glutamyl ligase |

| | | | |
|---|---|---|---|
| AEM58447.1 | G0HSS1 | 1Z69[41] | methylenetetrahydromethanopterin reductase |
| AEM58448.1 | G0HSS2 | | |
| AEM58449.1 | G0HSS3 | 8SBG[42] | tryptophanases |
| AEM58450.1 | G0HSS4 | 6F5Z[43] | methyltransferase |
| AEM58451.1 | G0HSS5 | | |
| AEM58452.1 | G0HSS6 | 3NBX[44] | Regulatory ATPase variant A - lysine decarboxylase |
| AEM58453.1 | G0HSS7 | | |
| AEM58454.1 | G0HSS8 | 6G49[45] | |
